# Vertebrate vision is ancestrally based on competing cone circuits

**DOI:** 10.1101/2024.11.19.624320

**Authors:** Chiara Fornetto, Thomas Euler, Tom Baden

## Abstract

Vision first evolved in the water, where light becomes increasingly monochromatic with viewing distance. The presence of spectrally broad (‘white’) light is therefore the exclusive remit of the visual foreground. However, if and how aquatic visual systems exploit this ‘white effect’ as an inductive bias, for example to judge distance, remains unknown.

By combining two-photon imaging with hyperspectral stimulation, genetic cone-type ablation, and behaviour, we here show that zebrafish suppress neural responses to the visual background by contrasting ‘greyscale’ and ‘colour’ circuits that emerge at the first synapse of vision. To do so, zebrafish use an early retinal architecture that fundamentally differs from that of mammals: Rather than combining cone signals to drive the retinal output leading to behaviour, zebrafish vision is built around competing ancestral cone systems: Red/UV versus green/blue. Of these, the non-opponent red and UV cones, which are retained in mammals, are necessary and sufficient for vision. By contrast, the colour opponent green and blue cones, which are lost in mammals, form a net-suppressive ‘auxiliary’ system that shape the ‘core’ drive from red and UV cones.

Our insights challenge the long-held notions that cones act in concert to drive visual behaviour, and that their spectral diversity primarily serves colour vision. Instead, we posit that vertebrate vision is ancestrally built upon opposing cone systems that emerged to exploit the strong spectral interactions of light with water. This alternative view points at terrestrialisation, not nocturnalisation, as the leading driver for visual circuit reorganisation in mammals.

## INTRODUCTION AND RESULTS

In image forming vision, information about two dimensions of space is hardwired into the retinotopic organisation of eyes^1^. By contrast, information about distance must be inferred. Terrestrial tetrapods including humans address this problem computationally, through a combination of inner retinal object-segmentation circuits^2–5^, stereoscopy^6,7^, and cognitive strategies such as the use of pictorial cues^8^. However, image forming eyes first evolved in the water^9^, which offers a much simpler cue for distance estimation: ‘Whiteness’ (Fig. 1a).

**Figure 1.**
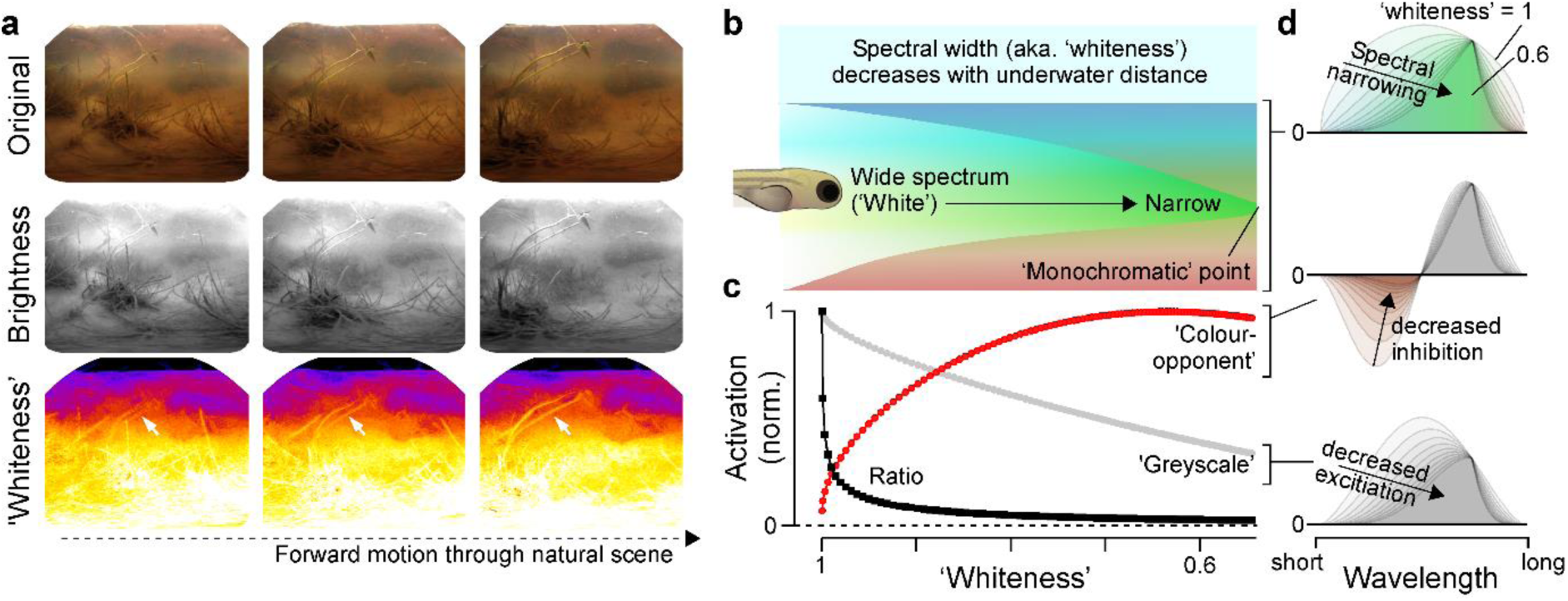
‘Whiteness’ decreases with underwater distance. **a**, Illustration of how spectrally broad (’white’) light fades with underwater distance, based on three subsequent RGB-colour frames (top) converted for their ‘brightness’ (middle) and ‘whiteness’ (bottom) content (Methods). Video data taken from zebrafish natural habitat, with the camera steadily moving forward through underwater vegetation^24^ (see also Supplemental Video V1). **b-d**, Schematic illustration of this ‘white-effect’ (b), and how idealised ‘colour-opponent’ and ‘greyscale’ detectors respond to a corresponding change in spectral width (c,d), as indicated (Methods).

Strong scattering and absorption of light in the water result in a rapid narrowing of spectral content with increasing viewing distance^10^ (Fig. 1b). The presence of spectrally broad light (i.e. ‘white’) is therefore the exclusive remit of the foreground (Fig. 1a,b). Aquatic visual systems could exploit this ‘white effect’, for example, by biasing retinal light responses towards spectrally broad input. Such a bias in the retinal output to the brain would almost inevitably lead to a foreground bias in visual behaviour. However, if and how aquatic visual systems use this inductive bias remains unknown.

In principle, a retinal white bias could be implemented by comparing achromatic (‘greyscale’) and chromatic (‘colour’) circuits^11,12^, because these tend to respond in opposite directions to a change in spectral width (Fig. 1b-d). Greyscale circuits (non-opponent) become progressively less active as the spectrum narrows (Fig. 1d, bottom) but colour circuits (opponent) become more active. This is because the excitatory and inhibitory inputs to colour opponent circuits cancel under spectrally broad light^13^, and this cancellation wanes as the spectrum narrows towards its monochromatic point (Fig. 1d, middle).

Any concurrent decrease in brightness and/or achromatic contrast^10^ then accentuates these spectral differences: Both strongly deteriorate greyscale signals, while colour circuits are largely invariant to these orthogonal axes in stimulus space.

All required components for contrasting greyscale and colour circuits are present in the larval zebrafish retina^13,14^. Like many fish, amphibians, reptiles and birds^9,15,16^, this diurnal surface-dwelling teleost retains all four ancestral cone-photoreceptor types (red, green, blue, UV; for definitions see Refs^9,16^). In the live eye, zebrafish red and UV cones are non-opponent, but green and blue cones are opponent due to outer retinal circuit interactions^13^. Accordingly, the four ancestral cone types of the zebrafish eye can be split into two ‘greyscale’ (red/UV) and two ‘colour’ channels (green/blue). Net antagonistic wiring of red/UV versus green/blue cone driven circuits would therefore lead to a white bias in the retinal output, which in turn would mean that the visual information available for behaviour is dominated by features from the (spectrally broader) foreground.

Here, we present direct evidence in support of this hypothesis. First, using two-photon imaging we demonstrate that zebrafish vision is profoundly white-biased. Second, using genetic ablation of individual and combinations of cone types, we show that this white bias emerges from the systematic contrasting of red/UV versus green/blue circuits. Specifically, we show that red and UV cones are necessary and sufficient for spatiotemporal vision, while green and blue cones are not necessary nor sufficient for vision and instead suppress red/UV circuits. Third, we show that the green and blue systems act in mutual opposition. Fourth, we confirm our results at the level of three ancient and highly conserved visual behaviours: Spontaneous swimming in the presence and absence of light, phototaxis, and the optomotor reflex.

Together, our results challenge the textbook notion that vertebrate photoreceptors act in concert to drive visual behaviour, while their opposition serves colour vision^12,17^. Instead, we propose that ancestral cone circuits are fundamentally wired to compete^9^. This alternative view on the retina’s originally aquatic circuit architecture opens new insights into the evolution of vision, and points at terrestrialisation^18^, not nocturnalisation^19^, as the leading driver of visual circuit reorganisation in early *Synapsida*, the ancestors of all mammals^20–22^.

### Zebrafish vision is white-biased

To assess how zebrafish process spectral content in spatiotemporal visual stimuli, we used two-photon imaging of light-evoked neural activity in the tectum, the principal retinorecipient area of the teleost brain (Fig. 2a,b). Stimuli were projected onto a lateral screen using a custom ‘four-colour hyperspectral’ stimulator^23^, with wavelengths approximating the spectral sensitivity peaks of the cones *in-vivo*^13^ (from ‘red’ to ‘UV’: 587, 470, 422 and 373 nm, respectively). Relative stimulus intensities were adjusted to follow the natural spectral distribution of daylight in the zebrafish habitat^24^ (from ‘red’ to ‘UV’: 1200, 600, 300 and 150 μW, respectively, Fig. S1a-c). Accordingly, concurrent activation of all four spectral channels approximated ‘zebrafish-white’ (achromatic), while selective activation of subsets could be used to present various ‘coloured’ (chromatic) stimuli.

**Figure 2.**
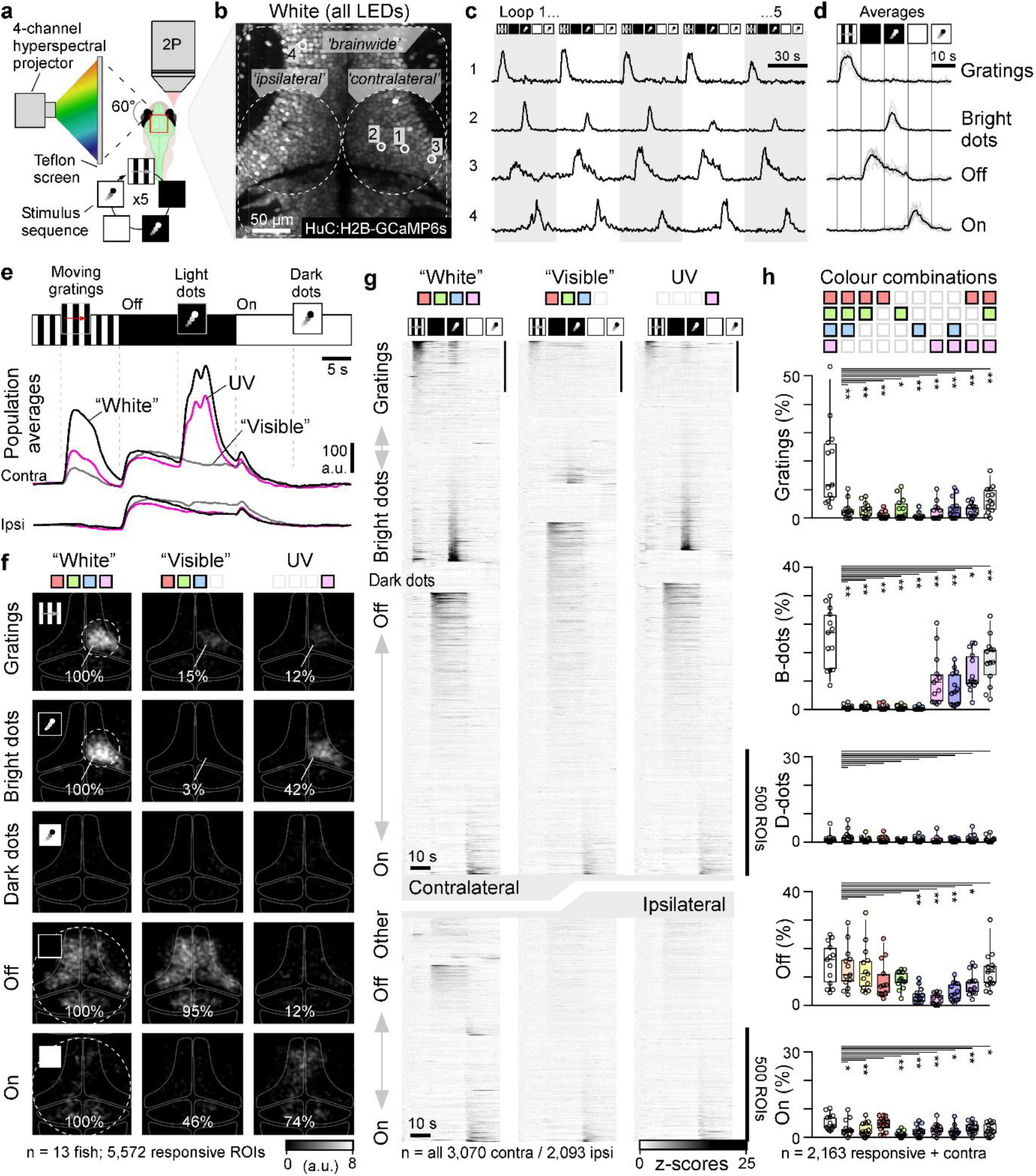
Zebrafish vision is ‘white biased’. **a**, Experimental setup for two-photon imaging of tectal neurons in larval zebrafish^68^ during presentation of ‘4-colour spatial stimuli’^23^. **b-d**, Representative scan with annotated brain regions and example ROIs (b) and their single trial (c) and averaged (d) responses to a battery of widefield and spatiotemporal stimuli as shown. **e**, Average responses to the same stimulus sequence of all contra-(top) and ipsilateral ROIs (bottom). ROIs from 13 fish when presented in ‘white’ (black line), ‘visible light’ (grey line), and UV (pink line). **f**, Spatial distribution of all analysed responses within the brain, as annotated. Percentages indicate the average abundance of different response types relative to ‘white’ (=100%). **g**, Overview of all contralateral (top) and ipsilateral (bottom) responses to white, visible and UV light, sorted by response type (see also Supplemental Video V2). **h**, Statistical evaluation of the number of responses to each stimulus and colour combination as annotated. Shown for each fish (n = 13) are numbers of ROIs that responded to a given stimulus as a percentage of all contralateral ROIs in a fish that passed a minimum response threshold (Methods) for any stimulus. Statistical comparisons against ‘white’ (1^st^ column) based on 1-tailed paired Wilcoxon Rank test with Bonferroni correction for multiple comparisons (*p <0.05; **p < 0.01; no asterisk: p>0.05). For full statistics, see Supplemental table T1.

Throughout, we focussed on three universal aspects of visual circuit computation:

i. Light detection (widefield On/Off), linked e.g. to phototaxis^25,26^ and visuomotor reflexes^27^.
ii. Widefield motion (large moving gratings), linked e.g. to optomotor/optokinetic reflexes^28^.
iii. Local object motion (small moving dots), linked e.g. to visual prey capture^29–31^.

Of these, the detection of light or its absence is principally possible with any photosensitive system, including extraocular, while the latter two require retinal processing (hereafter: ‘spatiotemporal vision’). To probe the above-described stimulus space, we used a stimulus sequence comprising widefield moving gratings as well as small light and dark moving dots superimposed on a dark and bright background, respectively (Fig. 2c-e). When presented in ‘zebrafish white’, individual tectal neurons responded selectively to specific aspects of this stimulus sequence over repeated trials (Fig 2c,d): Moving gratings and bright moving dots on a dark background (hereafter: ‘bright dots’) elicited large numbers of responses, while dark dots on a bright background (hereafter: ‘dark dots’) triggered few responses, if any. Together, responses to spatiotemporally patterned stimuli were consistently restricted to the retinotopically aligned base of the tectum, in line with direct drive from the stimulated eye (Fig. 2e,f, cf. Fig. 2a,b). By contrast, On/Off responses (i.e. light detection) were routinely observed all over the brain, with no obvious contralateral or retinotopic bias.

Next, we repeated this experiment using various shades of ‘coloured’ light (Fig. 2e-h). This revealed a striking white-bias in the representation of spatiotemporal stimuli, but not in basic light detection. For example, ‘visible light’ (RGB), lacking the ultraviolet (UV) component, was far less effective at eliciting spatiotemporal pattern responses compared to white (i.e. RGBU; cf. Fig. S1a-f): Omission of UV, which accounts for less than 7% of the total stimulus power in white, resulted in an 85% loss of grating responses, and 97% of bright dot responses (i.e. white-corrected efficiencies of 15% and 3%, respectively). Conversely, activating the UV-channel alone yielded a white-corrected efficiency of only 12% for gratings, and 42% for bright dots (Figs. 2e-h).

Other wavelength combinations yielded comparable results (Fig. 2h, Fig. S1f): Relative to white, the preponderance of both grating and bright dot responses was strongly and significantly reduced in all non-white stimulus conditions. For gratings, the most effective non-white stimulus was ‘RGU’, with a white-corrected mean efficiency of 36% despite a light power loss of less than 9%.

Spectrally narrow stimulation was even less effective, with ‘red’, ‘green’, or ‘blue’ channels (R, G, B) individually eliciting fewer than 5%, 17% and 3% grating responses relative to white. The most reliable bright-dot responses were also elicited by ‘white’ light, however unlike for gratings, and in line with previous work^24,29,32^, their presence or absence was in addition specifically related to presence or absence of UV light. The most effective non-white condition was again RGU with a white-corrected efficiency of 72%. RU and UV-only stimulation were second and third, with 55% and 42% efficiency, respectively. In the absence of UV light, independent of which other wavelengths were present, bright dot responses were elicited with a white-corrected efficiency of at most 5%. These results demonstrate that responses to spatiotemporally patterned stimuli are ‘white-biased’ far beyond what could be reasonably explained by stimulus intensity alone.

Unlike for responses to patterned stimuli, neither Off nor On-responses were specifically tuned to ‘white’. For example, in both cases the response to ‘red’-only (R) stimulation was statistically indistinguishable from the white response (Fig. 2h).

We conclude that in the larval zebrafish brain, the representation of spatiotemporally pattered stimuli such as drifting gratings or prey-like moving dots, but not of basic light-dark transitions, is under intimate control of spectrally competing circuits that suppress responses to non-white stimuli.

### Spatiotemporal vision does not require green or blue cones

To pinpoint the origin of the white bias, we generated transgenic zebrafish lines where each of the four types of cones could be selectively ablated^13^ and presented each with the same ‘white’ stimulus sequence. This directly confirmed our hypothesised^9^ circuit architecture: Ablation of red or UV cones, but not of green or blue cones, resulted in a profound loss in overall visual responsiveness (Fig. 3a). This loss was consistently characterised by a near-complete cessation of spatiotemporal pattern responses, but only moderate effects on Off and On responses (Fig. 3a-I, Fig. S2).

**Figure 3.**
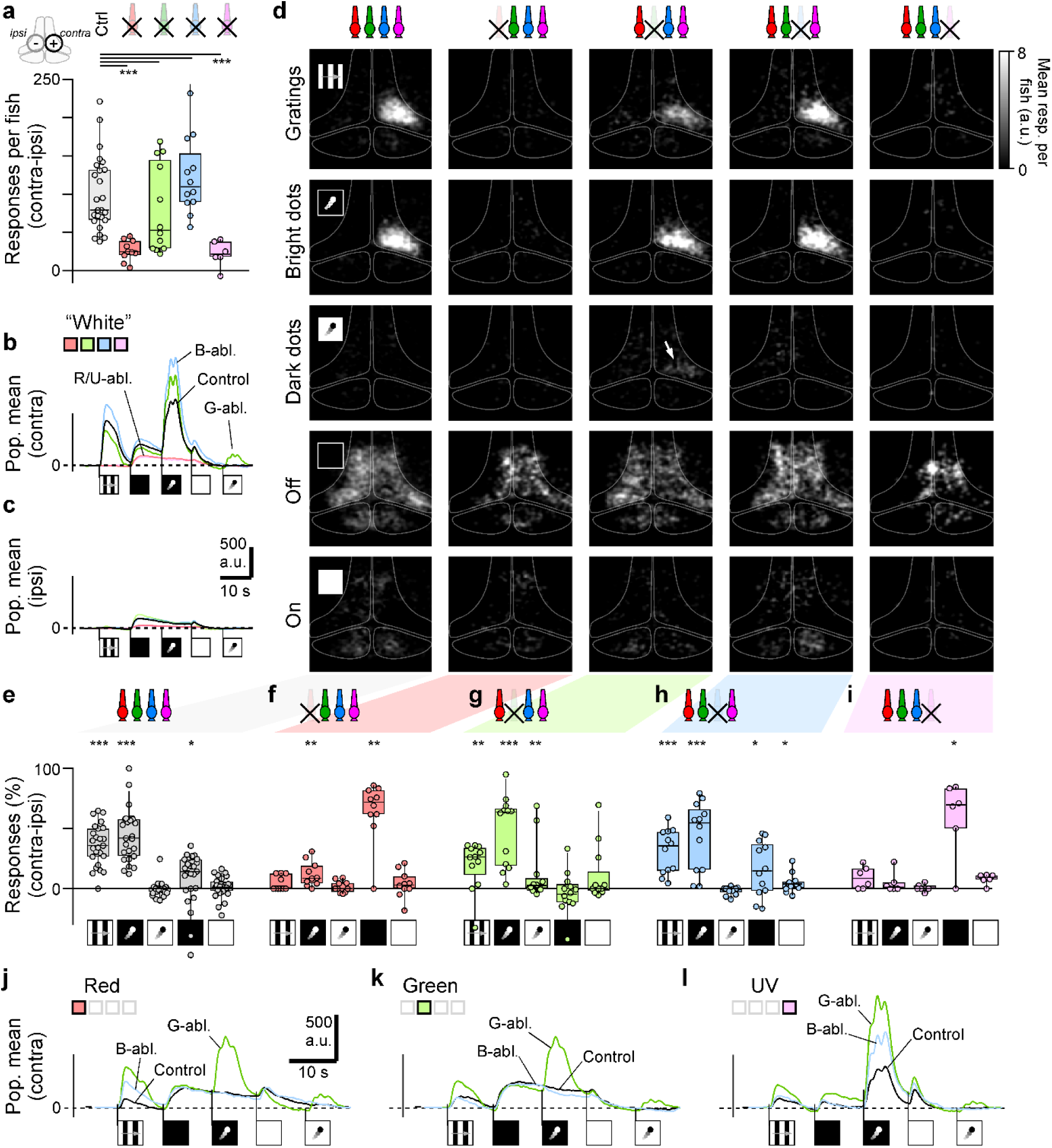
Spatiotemporal vision does not require green or blue cones. **a**, Number of contralateral responsive ROIs per fish that passed our minimum response criterion (Methods) during presentation of our battery of spatiotemporal stimuli (cf. Fig. 2) presented in ‘white’, minus the corresponding number of ipsilateral responses. From left: shown are responses for control animals and following the genetic ablation of red, green, blue and UV cones. Note that unlike for the ‘full colour’ data shown in Fig. 2, control data shown is based on a larger sample of n = 25 fish. Statistics based on 2-tailed Wilcoxon Rank Sum Test with Bonferroni Correction for multiple comparisons. For details, see Supplemental Table T2. **b-d,** as Fig. 2e,f, but only for ‘white’ responses of controls and following ablation of different cones, as indicated. The arrow in (d) highlights the unmasking of dark-dot responses in green ablated animals. For the underlying data, including numbers of fish and ROIs, see Fig. S2a-e. **e-i**, Statistical evaluation of contralateral responses per fish to different stimulus aspects as indicated, corrected for ipsilateral responses. Shown are the numbers of responsive ROIs per stimulus as a percentage of all responsive ROIs. Note that this normalisation overemphasises response numbers in poorly responsive animals (e.g following red/UV ablation). Uncorrected numbers of responses brain wide are shown in Fig. S2f-k. Statistical comparisons are made per stimulus condition comparing the numbers of contralateral versus ipsilateral responses, based on 1-tailed Wilcoxon Rank test (*p <0.05; **p < 0.01; ***p < 0.001; no asterisk: p>0.05). For full statistics, see Supplemental Table T2. **j-l**, as b, but instead of ‘white’, shown for ‘red’ (j), ‘green’ (k) and ‘UV’-stimulation (l).

First, ablation of red cones completely abolished grating responses but spared a small but significant number of bright dot responses (Fig. 3b-f). Second, UV cone ablation completely abolished both grating and dot responses (Fig. 3b-d,i). Third, both grating and bright dot responses persisted the ablation of green or of blue cones (Figs. 3b-d,g,h). Of these, response numbers were statistically indistinguishable from controls, except for green ablated grating responses, which were slightly but significantly reduced (p_B-dots_ = 0.24 (G) and 0.43 (B); p_Gratings_ = 0.013 (G) and 0.23 (B), 1-tailed Wilcoxon Rank Sum). Moreover, green cone ablation unmasked previously absent responses to dark dots (p_D-dots_ = 0.007 (G) and 0.043 (B)).

Ablation of green or blue cones also led to various types of increases in the overall ‘population response’ to spatiotemporally patterned stimuli, which considers both the number of responsive neurons and their individual response amplitudes (Fig. 3b,c). These effects tended to increase when using spectrally narrow light (Fig. 3j-l) instead of white. For example, blue cone ablation accentuated responses to red light gratings (Fig. 3j) and to UV bright dots (Fig. 3l). The same responses were also increased following green cone ablation, however in this case effects were more widespread and notably included the unmasking of dark dot responses at all tested wavelengths (Fig. 3j-l), as well as increases of spatiotemporal pattern responses under UV light (Fig. 3l), where green cones themselves are insensitive (Fig. S1e). The latter observation implies that green cones indirectly suppress UV responses, for example by interaction with blue cones (Fig. S1e).

Other than for the above-described effects on spatiotemporal pattern responses, the representation of Off and On stimuli mostly persisted individual cone-type ablations (Fig. 3b-i), and their contralateral bias tended to be weak (Off) or altogether absent (On) including in the control condition (Fig. 3d-h).

Nevertheless, the only significant contralateral changes were again in the case of green and blue cone ablation. The weak contralateral Off bias was selectively lost following green cone ablation, while blue cone ablation uniquely unmasked a small but significant contralateral bias in On-responses (Fig. 3h).

Together, the results from these ablation experiments strongly suggest that the processing of different types of basic visual stimuli is specifically linked to distinct subsets of cones^9^. In particular, the profound overall loss in responses to spatiotemporally patterned stimuli in the absence of red or UV cones indicates that these two cones are essential for ‘normal’ vision. Conversely, the persistence of spatiotemporal pattern responses following the ablation of green or blue cones, coupled to the observation that responsiveness to spatiotemporal patterns tended to increase, and novel response types could be unmasked, indicates that these cones normally suppress rather than drive the retinal output.

### Red and UV cones are necessary and sufficient for spatiotemporal vision

The inferred opposite roles of red/UV and green/blue circuits in vision were further directly supported by the results of double-ablation experiments: Concurrent ablation of red and UV cones resulted in a complete cessation of contralateral spatiotemporal pattern responses (Fig. 4a-f, Fig. S3). Conversely, spatiotemporal pattern responses persisted the concurrent ablation of green and blue cones (p_B-dots_ = 0.14; p_Gratings_ = 0. 14; Fig. 4g, Fig. S3). Moreover, as previously observed for green cone ablation (Fig. 3) dark dot responses were unmasked (p_D-dots_ = 0.023; Fig. 4b,g) and the contralateral bias for Off-responses was lost. However, concurrent ablation of green and blue cone types ameliorated the overall ‘inflation’ of spatiotemporal pattern responses that was observed following individual cone type ablations (Fig. 4b, j-l, cf. Fig. 3b, j-l).

**Figure 4.**
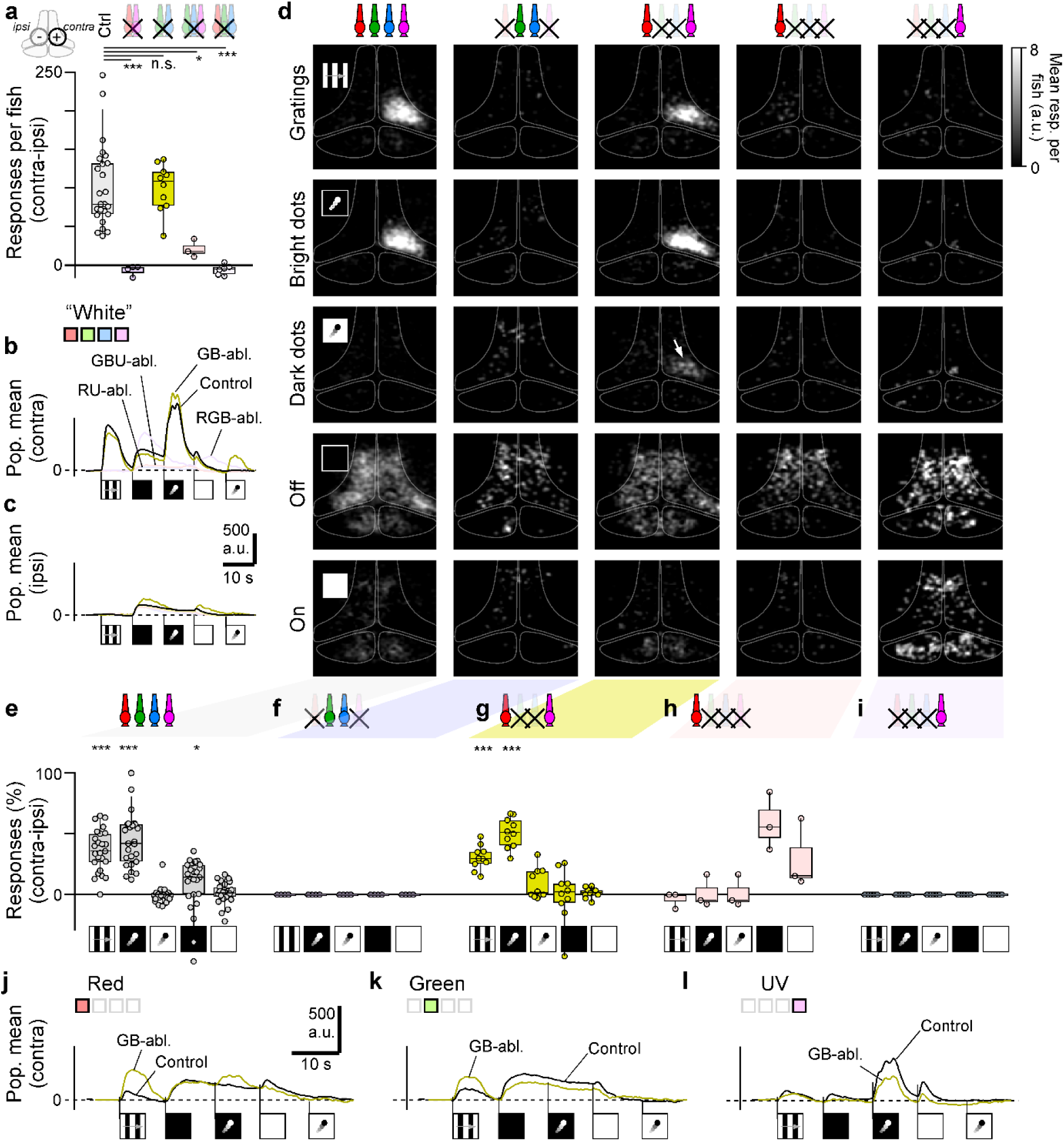
Red and UV cones are necessary and sufficient for spatiotemporal vision. **a-l,** as Fig. 3a-l, but here shown for a different set of cone-ablations. In each set of panels, from left: Control, red/UV double ablated, green/blue double ablated, green/blue/UV triple ablated, red/green/blue triple ablated. For statistics as shown in (e-l), see Supplemental Table T2. For clarity, panels j-l only show controls and green/blue double ablated data. Note that for convenience, the control data from Fig. 3 is repeated.

Next, even though concurrent ablation of red and UV cones approximately recapitulated their individual ablation phenotypes, the ‘normal’ functioning of the visual system was contingent on their joint presence: Triple ablations, leaving only red (Fig. 4h) or only UV cones in place (Fig. 4j), both resulted in a complete loss of pattern responses.

Together, the results from this series of ablation experiments strongly suggest that the joint presence of both red and UV cones is necessary and sufficient for normal spatiotemporal pattern vision. Conversely, the presence of green and blue cones appeared to be inessential for spatiotemporal vision (at least for the tested stimulus space), and their individual absence accentuated various aspects of visual responses.

The picture that emerges is one where the four ancestral types of cones of the vertebrate eye can be divided into two opposing systems: Red and UV cones drive vision, but green and blue cones suppress and thereby regulate visual circuits. Moreover, the ‘inflated’ response phenotypes following individual green or blue cone ablation, compared to green/blue double ablation, further implies that green and blue cones normally interact amongst themselves before imparting their regulatory functions upon red and UV cone circuits.

### Green and blue cones cause the white bias in zebrafish vision

Next, we systematically combined cone ablations and spectral stimulation to elucidate how green and blue cones regulate vision. To this end, we concurrently ablated green and blue cones and presented the same battery of spectral combinations that was previously used to probe controls (cf. Fig 2h, Fig. S1f). This revealed that for both gratings and bright dots, green/blue double ablated animals were systematically retuned relative to controls (Fig. 5a,b, Fig. S4a-d). In both cases, spectral responses were shifted towards longer wavelengths at the expense of shorter wavelengths. Moreover, the white bias was largely lost. For example, unlike in controls, grating responses to red-only stimulation were statistically indistinguishable from white stimulation in green-blue ablated animals (p = 0.11, Paired Wilcoxon Rank Test, 1-tailed). In fact, now the spectral tuning functions of gratings and Off-responses were essentially identical (Fig. 5a, cf. Fig. S4c). We conclude that the striking white bias in zebrafish vision (Fig. 2) stems from net-suppressive circuit interactions of green and blue cones on the retinal output.

**Figure 5.**
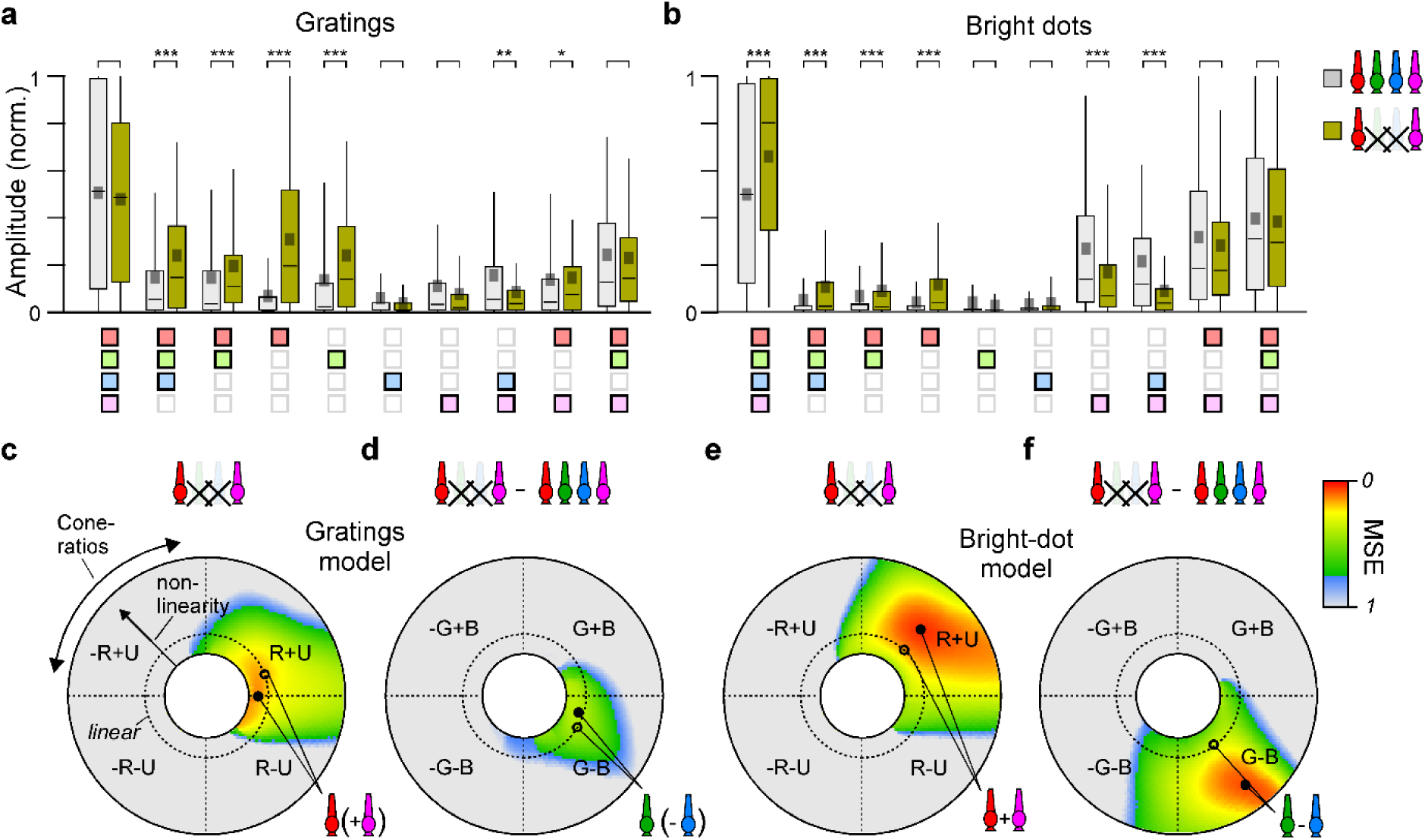
Green and blue cone circuits act in mutual competition. **a,b**, Normalised spectral response amplitude distributions of all contralateral ROIs that responded to gratings (a) or bright dots (b) in controls (grey) and in green/blue double ablated animals (yellow). For details on the normalisation procedure, see Methods. The full green/blue double ablated data across all ten spectral combinations is summarised in Fig. S4a, and amplitude distributions for dark dots as well as Off and On stimuli are shown in Fig. S4b-d, respectively. Statistical comparisons in (a,b) based on 1-tailed Wilcoxon Rank test with Bonferroni correction for multiple comparisons (*p <0.05; **p < 0.01; ***p < 0.001; no asterisk: p>0.05). For full statistics, see Supplemental Table T3. **c-f**, Results of a two-step cone-combinatorial model (Methods) that captures the spectral tunings of grating (c,d, cf. a) and bright dot responses (e-f, cf. b) using the spectral tunings of cones (cf. Fig. S1d,e). Heatmaps summarise the mean squared error (MSE) between a target tuning function and the tuning of each possible cone combination, as indicated. MSE normalised between 0 (perfect fit) and 1 (no fit). Distance from the centre denotes the linearity of the model from sublinear to supra-linear towards the outside (Methods). The dashed circle denotes the linear point, while closed and open circles denote the best fit overall and the best linear fit, respectively. Corresponding best fits are shown in Fig. S4e-h. Inferred cone ratios (2 s.f.) as follows: Gratings: 0.99R:-0.01U (linear: 0.75R:0.25U); 0.88G:-0.12B; Dots: 0.45R:0.55U; 0.42G:-0.58B.

### Green and blue cone circuits act in mutual competition

We next used the spectral tuning functions of control and green/blue double ablated animals to computationally infer the underlying cone contributions to grating and dot responses. For this, we used a two-step cone-combinatorial model^13,33^ (Fig. 5c-f). Data from green/blue double ablated animals was used to infer red versus UV cone weights (Fig. 5c,e), while the differences between controls and double ablated animals were used to infer green and blue cone weights (Fig. 5d,f). Together, this added further support to the notion that red and UV cones act in concert while green versus blue cones act in mutual opposition.

For gratings, spectral responses of green/blue ablated animals were best approximated by a sublinear drive from red cones alone (87.3% captured, Fig. 5c, Fig. S4e). Small additional contributions from UV cones achieved near equivalent and linear fits (82.7%, Fig. 5c, open symbol). Conversely, the spectral differences between double ablated and control responses were best explained by net-positive contributions from green cones but net negative contributions from blue cones (61.3%, Fig. 5d, Fig. S4g). Likewise, bright dot tuning functions were best explained by a net positive contribution from both red and UV cones (95.1%, Fig. 5e, Fig. S4f) but an opposing contribution from green minus blue cones (92.4%, Fig. 5f, Fig. S4h). Accordingly, both response types point at a single underlying circuit architecture: Joint drive from red and UV cones, and mutual opposing regulation from green and blue cones. The main difference between grating and bright dot circuits appeared to be that the former is red/green dominant and (sub)linear, while the latter is UV/blue dominant and supra-linear.

Beyond gratings and bright dots, the newly unmasked responses to dark dots also followed a bright-dot-like “U-shaped” spectral tuning curve, indicative of concomitant red plus UV cone drive (Fig. S4b). In this case, the sparsity of dark-dot responses in control animals precluded estimating relative green/blue contributions. However, the observation that dark dot responses were selectively unmasked following green cone ablation (Fig. 3g), but not blue (Fig. 3h), points to an unequal suppressive action of green versus blue systems in controls.

### Concurrent ablation of green and blue cones restores lost balance that follows single cone ablations

If green and blue cones act in mutual opposition to regulate red/UV-driven circuits, then selective ablation of only parts of such a regulatory system should result in a poorly balanced visual circuit. However, this balance should improve if the regulatory system is ablated in its entirety. For example, individual differences between animals might be expected to exacerbate if the green/blue system is damaged, but less so if it is altogether missing. This effect was in fact readily observed in the scattered response distributions of across green- or blue- single ablated animals (Fig. 3g,h) compared to controls (Fig. 3e), as well as the notably tighter distribution of responses across animals for green-blue double ablated animals (Fig. 4g). To quantify this effect, we computed the correlation of each animals’ response-distribution (i.e. the relative responsiveness to gratings, bright dots, dark dots, Off, and On) against the remainder of the population, such that 1 and 0 indicate perfectly homogeneous and random populations, respectively (Fig. 6a). This confirmed that both green and blue single-ablated animals were significantly more heterogeneous than controls, while green/blue double ablated animals were significantly more homogeneous.

**Figure 6.**
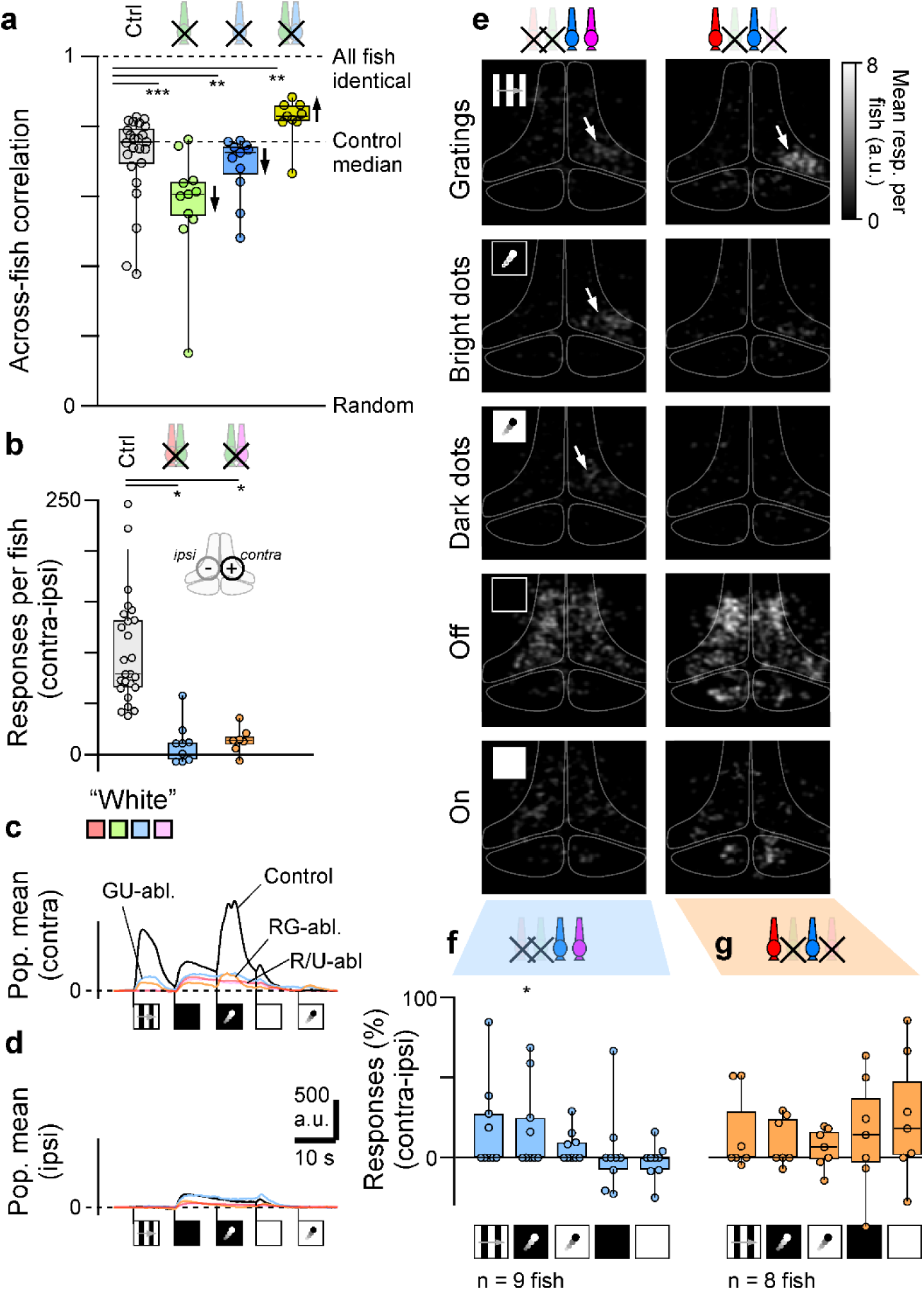
Restoration of lost balance by concurrent green/blue ablation. **a**, Response homogeneity (Method) across the population of control animals and following ablation of green, blue and green/blue cones, as indicated. Statistical evaluation based on 1-tailed Wilcoxon Rank test of each ablation condition relative to controls, with Bonferroni correction for multiple comparisons. pG = 0.000036, pB = 0.0004, pG/B = 0.0061. **b-g**, as Fig. 3a-I, but here shown for controls and two new ablation combinations: red/green (left), and green/UV (right). For full statistics, see Supplemental Table T2.

To further explore this idea, we carried out two additional sets of double-ablation experiments, however this time ‘half-ablating’ both the core and the regulatory systems. We reasoned that ‘half-ablating’ the core system (i.e. red or UV, but not both) should disrupt spatiotemporal pattern responses, while ‘half-ablating’ the regulatory system (here: green but not blue) should then part-counteract this disruption. This is indeed what we found. Both red/green and UV/green double-ablated animals were less responsive than controls, but more responsive than red or UV single-ablated animals (Fig. 6b-e, cf. Fig. 3, Fig. S5). Moreover, strikingly, these ‘half-ablated’ animals were exceptionally heterogeneous: Following either ablation pair, approximately half of animals were entirely unresponsive to any spatiotemporal patterns, while the other half displayed response levels that were well within the normal distribution of controls (Fig. 6f,g, cf. Fig. 3e). These results underscore the antagonistic balance within the green/blue system, such that their experimental disruption can lead to a wide spectrum of phenotypic diversity.

### Cone ablations lead to type-specific deficits in spontaneous swimming behaviour

We next explored if and how different cone circuits can be linked to behaviour. We began with a characterisation of spontaneous behaviour by filming free-swimming individuals in constant light or constant dark (Fig. 7, Fig. S6). This revealed that all tested ablations led to cone type specific behavioural deficits in spontaneous swimming.

**Figure 7.**
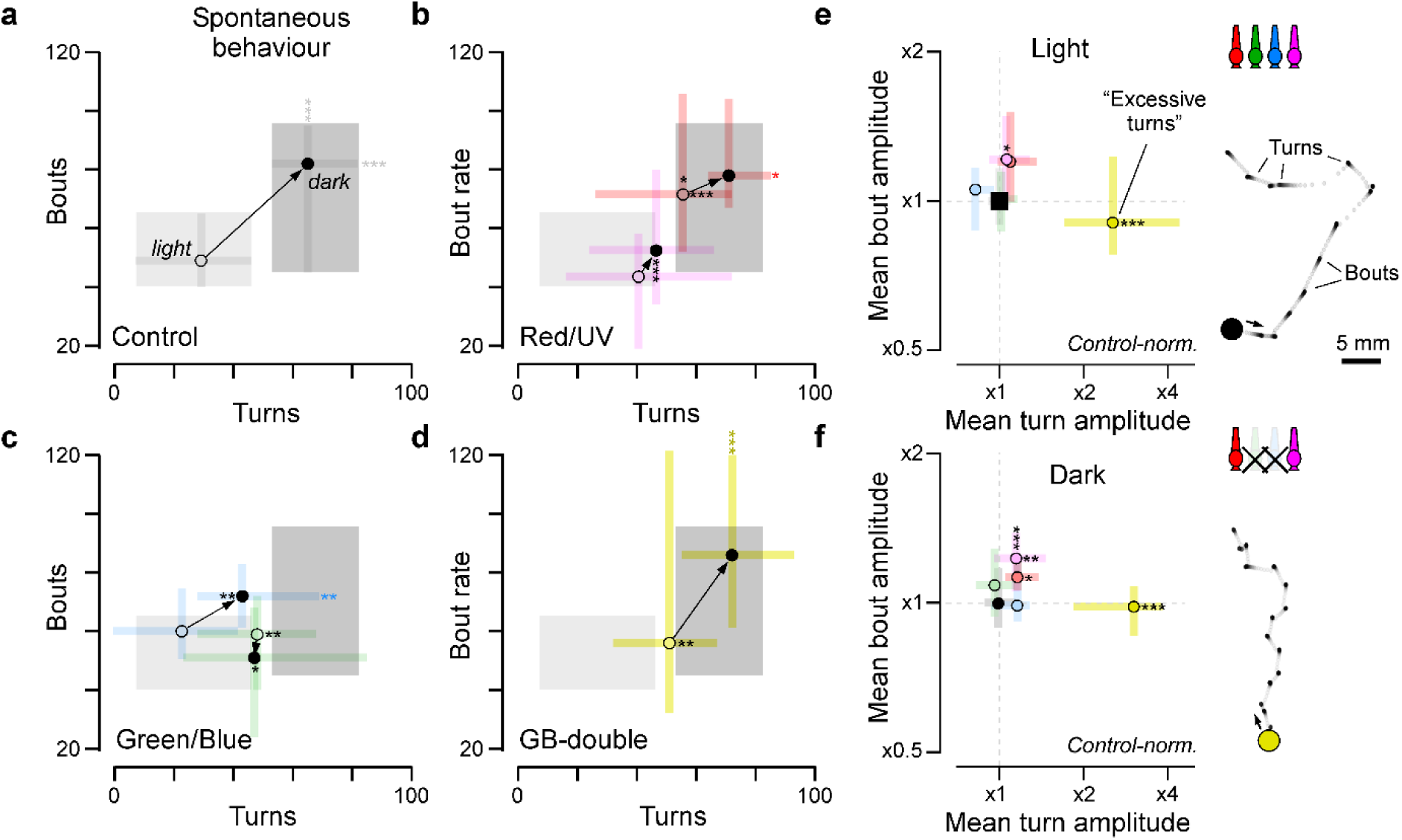
Cone type-specific deficits in spontaneous swimming behaviour. **a-d**, Spontaneous movement statistics (Methods) of free-swimming fish in continuous light (open symbols) or continuous dark (closed symbols), as indicated. Shown are controls (a), red and UV ablated (b), green and blue ablated (c) and green/blue double ablated animals (d). Shadings denote the interquartile ranges of light and dark behaviours in controls. For clarity, only population medians and interquartile range are plotted. Two sets of comparisons were performed, using 1-tailed Wilcoxon Rank tests: ‘within-metric’, comparing control versus each ablation condition (black asterisks placed within the corresponding range bars), and ‘within-condition’ comparing light versus dark (coloured asterisks placed outside of the error bars). For full statistics see Supplemental Table T4. **e,f**, Quantification of each populations’ median bout and turn amplitudes, normalised to controls (=1). For full statistics see Supplemental Table T5. Statistical evaluation based on 1-tailed Wilcoxon Rank tests with Bonferroni correction.

As shown previously^34^, control animals exhibited notably more swim bouts and spontaneous turns in the dark compared to the light (Fig. 7a). By contrast, this light dependent phenotype was largely lost following the ablation of red or UV cones (Fig. 7b). Instead, UV-ablated animals behaved as if always in the light, while red-ablated animals behaved as if always in the dark. These results support the notion that the concurrent presence of both cones is required for normal visual behaviour, and further hint that in the intact system, their relative activations contribute to determining behavioural state.

Next, unlike for red or UV, blue ablated animals behaved normally in the light but failed to fully reach control-like activity levels in the dark. Conversely, green ablated animals always exhibited intermediate activity levels. (Fig. 7c). Green and blue phenotypes were however ‘rescued’ in green/blue double ablated animals which, despite an elevated turn rate in the light, exhibited the most control-like behavioural performance overall amongst all tested ablations (Fig. 7d). These results add direct support to the notion that green/blue cones compete to regulate rather than drive vision.

However, despite their approximately control-like spontaneous activity levels, green/blue double ablated did exhibit another, unique behavioural deficit: They ‘over-turned’ (Fig. 7e,f). Independent of illumination, these animals exhibited more than two-fold larger mean turn amplitudes compared to any other tested population. It appears that in the absence of the ‘regulatory’ green/blue system, zebrafish struggle to reliably swim in a straight line.

### Cone-type specific optomotor deficits

We next tested the effects of cone ablations on optomotor performance, a probably ancestral reflex that helps animals to stabilise their body position relative to the visual environment^28^. We used a closed-loop setup^35^ where the orientation of sideways drifting gratings constantly updated to align with the primary body axis. In this environment, free-swimming control fish perpetually turn towards the widefield drift which leads to ‘circling behaviour’ (Fig. 8a). By contrast, ablation of different cone types led to various types of behavioural deficits (Fig. 8b-f).

**Figure 8.**
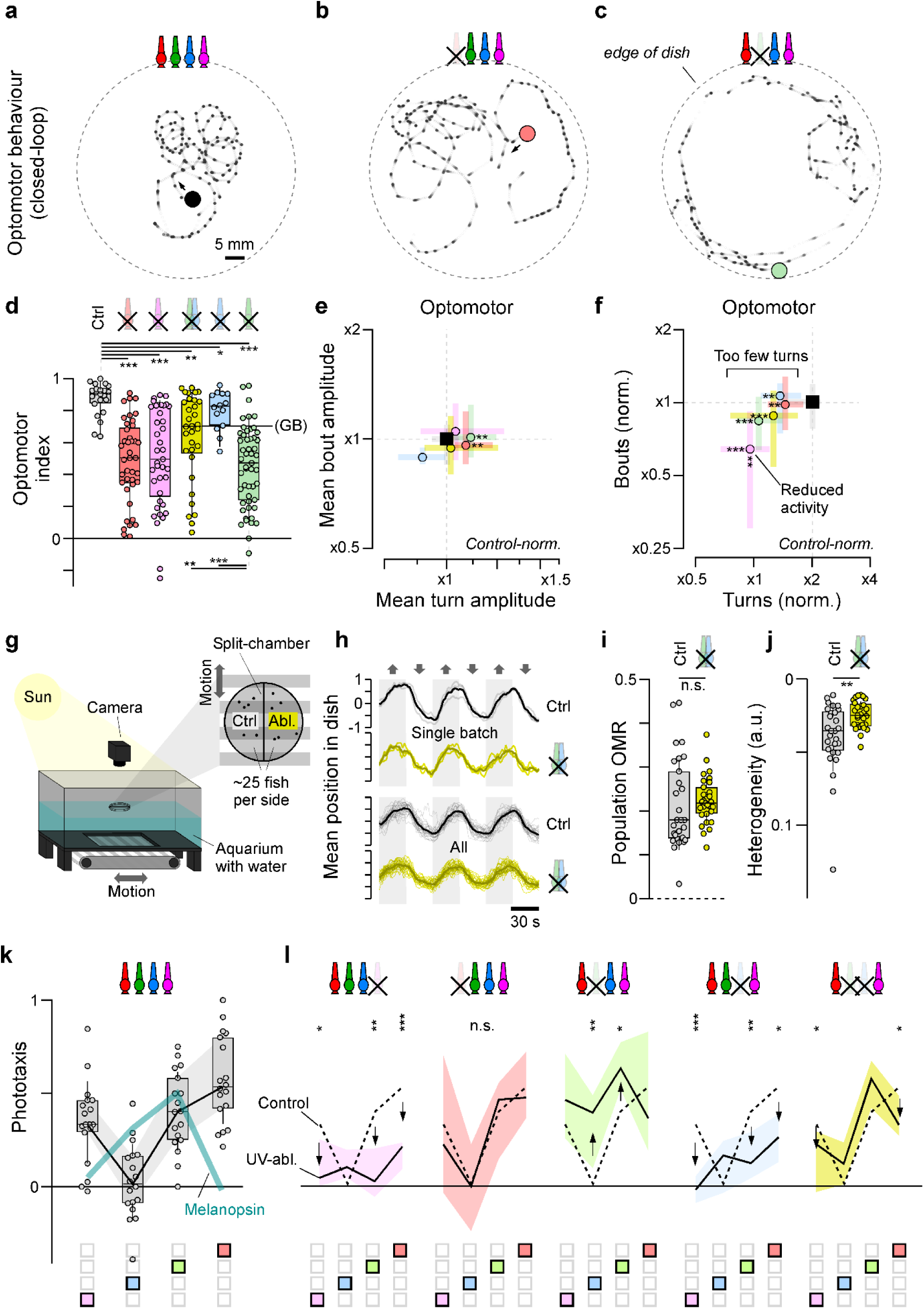
Cone-type specific deficits in optomotor behaviour and phototaxis. **a-c,** Example movement trajectories in closed loop optomotor assay (based on Ref^35^) for a control animal (a), a red-ablated animal (b) and a green-ablated animal (c). **d**, Quantification of optomotor performance (Methods) in controls and following different cone ablations as shown. Statistical evaluation based on 1-tailed Wilcoxon Rank Sum test with Bonferroni correction for multiple comparisons. Two sets of pairs were evaluated: Controls versus each ablation condition (top), and, separately, green/blue double ablated animals versus blue and green single ablated animals (bottom) (*p <0.05; **p < 0.01; ***p < 0.001; no asterisk: p>0.05, for full statistics see Supplemental Table T6). **e,f**, as Fig. 7e and Fig. S6a, respectively, but here shown for movement statistics during optomotor stimulation. **g**, Schematic of outdoor-optomotor arena (Methods). **h**, Average position of all fish per side over time as indicated. Top: three repeats of one experimental batch; bottom: all experiments (n = 29). For details, see Supplemental Figure S7. **i,j**, Quantification of mean optomotor performance per batch (i) and their heterogeneity across experiments (j). **k**, Spectral dependence of phototaxis behaviour in controls (grey) as indicated, with power-adjusted spectral tuning of melanopsin superimposed, assuming peak absorption (λ_max_) at 480 nm. **l**, Corresponding spectral dependences of phototaxis behaviour following the ablation of different cone types as indicated. For simplicity, only medians and interquartile ranges are shown, with the dotted line indicating control performance. Statistical evaluations of each spectral position against controls based on 1-tailed Wilcoxon Rank Sum tests with Bonferroni correction for multiple comparisons (*p <0.05; **p < 0.01; ***p < 0.001; no asterisk: p>0.05, for full statistics see Supplemental Table T7).

In general, deficits in the central representation of grating responses (Fig. 3,4) were a strong predictor of optomotor performance: Red, green and UV ablated animals exhibited poor optomotor responses, in line with the complete (red/UV, Fig. 3d,f,i) or moderate (green, Fig. 3d,g) loss of grating responses in the brain. By contrast, blue ablated animals displayed no significant defects, in line with their intact central representation of moving gratings (Fig. 3d,h). Moreover, as for spontaneous swimming (Fig. 7c,d), double ablation of green/blue cones partially rescued the green single-ablation phenotype. This behavioural pattern also implies that in the intact system, green circuits normally suppress blue circuits, which in turn suppress optomotor circuits: The only experimental difference between green and green/blue double ablated animals is the presence of blue cones. Accordingly, and perhaps counterintuitively, the poor behavioural performance of green ablated animals must be attributed to a disinhibited net-suppressive action of blue cones.

Next, the presence of high contrast moving gratings also largely overrode the spontaneous overturning phenotype of green/blue double ablated animals (Fig. 8e, cf. Fig. 7e,f). In fact, all tested ablation variants turn less frequently than controls (Fig. 8f), but all but blue also part-compensated for this reduction by increasing turn amplitudes (Fig. 8e).

Moreover, UV-ablated animals alone also exhibited reduced bout rates (Fig. 8f). The latter observation further hints that beyond visual deficits, the poor optomotor performance of red and UV ablated animals may also be part-related to systematic changes in behavioural state: UV-ablation tended to reduce behavioural activity, while red-ablation tended to result in hyperactivity (Figs. 7b,e,f, 8e,f).

### Green-blue ablation reduces optomotor variability under natural daylight

Encouraged by the close alignment between the representation of moving gratings in the brain (Figs. 3,4) and optomotor performance under artificial but controlled indoor conditions (Fig. 8a-f), we next wondered how controls and green-blue double ablated animals would perform under more natural, outdoor conditions. For this, we devised an outdoors optomotor arena based on an open-top water-filled aquarium suspended above a motorised conveyor belt for stimulus presentation (Fig. 8g, Methods). The arena was placed outdoors into the midday sun. We then suspended a split-chamber with ∼25 controls and green/blue double-ablated fish each on either side on the water surface to internally control for natural changes in lighting conditions over the course of experiments (clouds, shadows). We then moved the conveyor belt back and forth and quantified the mean position of all fish per side over time (Fig. 8h). This revealed robust and statistically indistinguishable optomotor performance across both populations of animals (Fig. 8i), both when the same animals were tested repeatedly in close succession (8h, top), and when comparing across different animal batches, experimental days, and water conditions (8h, bottom, cf. Supplemental Figure S7, Methods). Despite this, however, green-blue double ablated animals behaved significantly more internally consistent compared to controls (Fig. 8j). This reduced behavioural variability following the loss of green/blue cones, apparently without deteriorating overall performance, strikingly echoes their correspondingly reduced heterogeneity at the level of brain responses to spatiotemporally patterned stimuli (Fig. 6a).

### Cone regulation extends to intrinsically photosensitive circuits

Beyond rods and cones, vertebrate eyes also comprise intrinsically photosensitive circuits^36^ that mostly serve ‘non-image forming’ visual functions including the control of phototaxis^26^. This ancestral and probably universal behaviour allows animals to use information about the intensity and spectrum of light to seek out optimal places in their environment^37^. In larval zebrafish, experimental ablation of intrinsically photosensitive retinal ganglion cells (ipRGCs) disrupts this behaviour^26^. Nevertheless, ipRGCs cells do receive upstream synaptic inputs^36^ and might therefore be subject to the same types of cone-regulatory influences that also impinge on red/UV cone driven circuits.

To test this idea, we first determined the spectral dependence of phototactic behaviour in controls. Melanopsin, the photopigment expressed in ipRGCs, is ‘green’ sensitive (λ_max_ ∼480 nm)^36^, so in the absence of additional inputs from cones, phototaxis should be best driven by green light. However, this was not the case. Instead, phototaxis was spectrally biphasic and was best driven by UV or red light (Fig. 8k). We therefore next ablated different cones to probe which types contribute to phototactic behaviour, and how (Fig. 8l).

First, UV cone ablation led to highly significant reductions in phototactic performance, including at long wavelengths where UV cones are insensitive. Accordingly, while the loss of the UV response points at a direct role of UV cones in driving phototaxis, the more general loss of responsiveness across all wavelengths rather argues for an effect on behavioural state. Second, red cone ablation had no significant effect on phototaxis. This indicates that it is principally possible to robustly drive ‘basic’ visual behaviour without the use of red cones, despite their apparently indispensable roles in spatiotemporal vision (Figs. 2-7, 8a-f). Third, green cone ablation strongly accentuated phototactic behaviour, and moreover shifted the overall spectral tuning to more closely align with a potential joint drive from UV cones and melanopsin alone. This implies that green cones normally suppress phototactic behaviour. Fourth, and opposite to green, blue cone ablation reduced phototactic performance. Accordingly, blue and green cone circuits again acted in mutual opposition. However, importantly, in the case of phototaxis the roles of green versus blue cones were inversed, with green providing the principal suppression, and blue counteracting this effect. Finally, as already observed for both spontaneous swimming (Fig. 7d) and optomotor performance (Fig. 8d), concurrent ablation of both green and blue cones again rescued both green and blue single-ablation phenotypes to near control levels (Fig. 8l).

### A new view on the ancestral organisation of cone circuits for vertebrate vision

Our results challenge the ‘classical view’ on the functional organisation of ancestral cone circuits in the vertebrate retina (Fig. 9a). In this view, visuo-behavioural circuits are dominated by greyscale signals that emerge from a weighted sum across cones^17,38^. Antagonistic cone contributions, where present, are solely attributed to colour vision^12,38^. However, this view cannot explain:

i. The complete loss of responses to the tested spatiotemporal patterns following the ablation of red and UV cones (Figs. 3,4), because blue and green cones should continue to drive relatively normal vision in their absence.
ii. The widespread white bias of zebrafish vision (Fig. 1,2), the loss of this bias following the ablation of green/blue cones (Fig. 5), or the concurrent accentuation of spatiotemporal pattern responses (Figs. 3,4), because explaining any of these results requires a net-negative contribution of green and blue cones to vision.
iii. The ‘rescuing’ of green and blue cone single ablation phenotypes upon the concurrent ablation of both cones (brain functions: Figs. 3-6; behaviours: Figs. 7,8), because in the classical view, ablating cones cannot improve performance.

**Figure 9.**
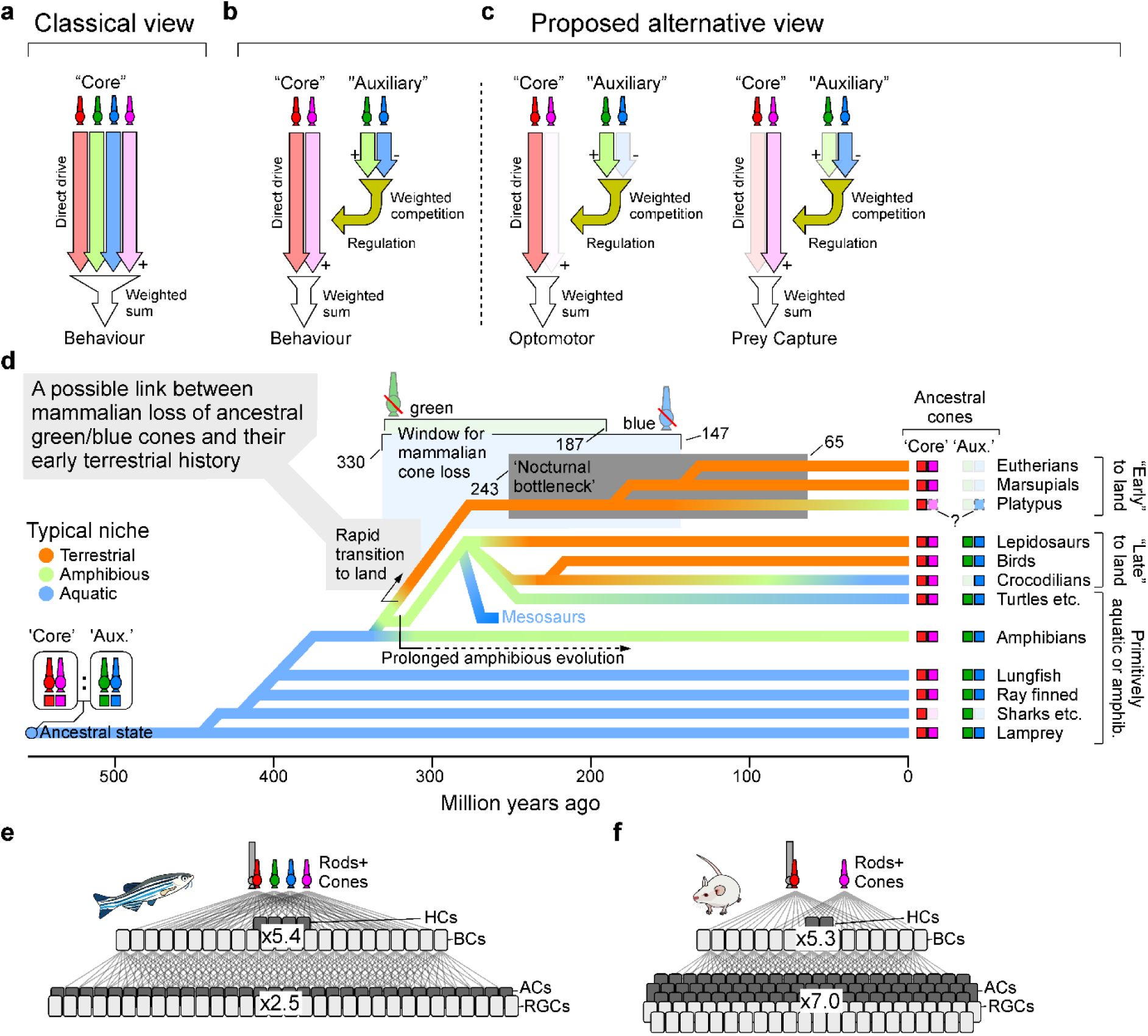
A new view on the organisation of cone circuits for vertebrate vision. **a,** ‘Classical’ view of cone signal cooperating to jointly drive behavioural circuits. **b**, Proposed alternative circuit architecture, where red and UV cones alone represent retinal ‘core’ circuits that jointly drives all visual behaviour. By contrast, green and blue circuits form an auxiliary circuit that acts in mutual opposition to suppress and thereby regulate the core drive. **c**, Suggested weight adjustment within this architecture for different subset of visuo-behavioural circuits. **d**, Approximate phylogenetic history of extant vertebrates with inferred ancestral patterns of territorialisation coded in colour, from the obligatory aquatic beginnings (blue) via amphibious lifestyles (green) to full terrestrialisation (orange). For background and key references, see Supplemental Discussion. Note that terrestrialisation probably occurred rapidly in the lineage that led to modern day mammals, but much more gradually so in the lineage that led to modern day reptiles including birds. This pattern correlates the widespread retention of the ancestral auxiliary cones in most non-mammalian tetrapods, but their systematic loss in probably all mammals (for platypus, see supplemental discussion). **e,f**, Schematic illustration of retinal neuron-type numbers in zebrafish (e) and mouse (f), from rods and cones (top) to horizontal and bipolar cells (HC, BC, respectively), which in turn feed into amacrine cells (ACs) and retinal ganglion cells (RGCs). The superimposed numbers summarise the corresponding neuron-type divergence ratios across layers. For references, see main text.

Instead, our results strongly support an alternative model^9^ of the functional organisation of ancestral cone circuits in the vertebrate retina (Fig. 9b):

Vision is primarily driven by weighted contributions from ancestral red and UV cones alone. Rods, which are not yet mature in larval zebrafish^39^, probably extend this principal drive to low light conditions^9^. Intrinsically photosensitive circuits add further drive to a subset of circuits. Together, these four photoreceptor types form the ancestral ‘core’ of vision, and for normal function, their concurrent presence is probably non-optional. In agreement, nearly all extant vertebrates, including humans, retain all four systems in their eyes^9,20^.

Conversely, ancestral green and blue cones do not drive vision. Instead, they form an ‘auxiliary’ system tasked with regulating the core circuits. Within auxiliary circuits, green and blue cones interact in mutual opposition, and the net result of this interaction, including its sign, dictates the nature of the regulatory input to different core circuits (Fig. 9c). The presence of green and/or blue cones for robust vision is probably inessential, in line with the profusion of vertebrate lineages, including humans, that have independently lost them^9^.

### Vision in air

We have shown that in zebrafish, one central role of the auxiliary cone system it to spectrally suppress visual responses to the background. It seems reasonable to suggest that other fish could use the same strategy. However, this ‘foreground trick’ does not work on land due the greatly reduced scattering and absorption of light in air^40^. Above the water, a fish-like spectral strategy for distance estimation can therefore only operate over tens of meters to kilometres – a visual range of limited behavioural relevance for most terrestrial species. This raises the question if and how a fish-like cone auxiliary system can continue to support vision in air (Fig. 9d).

Most extant terrestrial vertebrates belong to one of three major tetrapod lineages: *Lissamphibia* (salamanders, newts and frogs), *Sauropsida* (reptiles including birds), and *Synapsida* (mammals). Of these, most descendants of *Lissamphibia* and *Sauropsida* retain the complete set of ancestral cones in their eyes to the present day^9,16^. By contrast, no extant mammal retains ancestral green cones, and the same is probably also the case for ancestral blue (for a note on platypus, the only suspected exception^41^, see supplementary discussion).

Accordingly, the complete auxiliary cone system might have been lost as early as ∼330 million years ago, immediately following the divergence of *Synapsida* and *Sauropsida* (and notably contrasting Walls’ nocturnal bottleneck theory which has dominated thinking on this point^19^, see supplemental discussion). This important moment in vertebrate evolutionary history is also marked by a second event: A rapid move to the land by the ancestors of all mammals, but a probably much more gradual and incomplete transition to land in early reptiles^18,42^ (Fig. 9d). It therefore seems reasonable to suggest that the retention of ancestral green and blue cones in *Lissamphibia* and *Sauropsida* was linked to the continued utility of this system for supporting underwater vision. In early reptiles, a probably more than 50 million years window of mostly amphibious lifestyle then probably permitted the gradual emergence of compromise solutions for vision that worked both in water and in air. Conversely, in the lineage that led to modern mammals, the ancestral blue/green system could have presented a competitive disadvantage for exclusive vision in air, for example because its net-inhibitory architecture limited overall signal to noise.

At the same time, the loss of access to a relatively simple spectral strategy for foreground enhancement should have also created powerful selection pressures in early synapsids to find new solutions for visual figure ground segmentation in air. In modern mammals, part of this need is thought to be met by inner retinal circuits^2,3,43^. In this regard, it is striking to note a radical shift in the numerical abundance of ‘early’ versus ‘late’ retinal circuitry when comparing fish and mammals.

For example, zebrafish feed the signals from 5 outer retinal photoreceptor types (rods and 4 cones^16^) into 4 horizontal^13^ and 23 bipolar cell types^33,44,45^, which in turn serve a ‘relatively modest^46^’ ∼32 retinal ganglion cell types^26^ and a probably similar or even smaller number of amacrine cells^47,48^ (Fig. 9e). By contrast, mice feed the signals from 3 photoreceptor types into 2 horizontal and 14 bipolar cell types^20,49,50^, which then however serve some ∼45 types of ganglion cells^51,52^, and ∼67 types of amacrine cells^53–55^ (Fig. 9f). Accordingly, despite near-identical cell-type divergence in the outer retina (zebrafish: ∼5.4x, mice ∼5.3x), zebrafish and mice differ drastically in their degree of inner retinal divergence (∼2.5x versus ∼7.0x, respectively). One interpretation of this mammalian explosion in late retinal circuitry is that in the absence of outer retinal regulatory circuits provided by the auxiliary cone system and their immediate downstream horizontal and bipolar cells, early synapsids compensated for this loss by the emergence of new inner retinal regulatory circuits. Modern birds and reptiles then appear to straddle these extremes, with ‘fish numbers’ of early retinal neurons, but ‘mammalian numbers’ of late retinal neurons^9,20,40,56,57^.

### Possible circuitry underlying the green/blue system’s net-suppressive wiring

The net-suppressive action of ancestral green and blue circuits, including their mutual antagonism, could be relatively easily achieved by known retinal circuit organisation^9,14,17,33,58^. In the outer retina, horizontal cells set up the cone opponencies in the first place^13^, while bipolar cells then pool the signals only from spectrally neighbouring cones^45^ (‘spectral block wiring^12^’) in a type-specific manner. In doing so, some but not all bipolar cells invert the intrinsic ‘Off’-polarity of cones to set up parallel On and Off pathways^58^ (‘pathway splitting^59^’). As a population, bipolar cells therefore represent a wide but highly systematic mixture of cone signals that exist in two polarities^24,33,48^. The sign-conserved (‘Off’) and sign-inverted (‘On’) cone mixtures are then mapped onto developmentally hardwired depths of the inner retina, resulting in a ‘spectral layering^9^’ that superimposes on the traditional division by polarity^58^. Moreover, the multistratified nature of bipolar cells in fish^45^ means that On and Off signals routinely coexist within common layers of the inner retina^48^. This key anatomical feature is shared with birds^57,60^, reptiles^61^ and amphibians^62^, all of which retain the complete ancestral cone complement. By contrast, multistratified bipolar cells are largely absent in mammals^58^.

In fish, the resultant inner retinal layering presents an ideal substrate for flexibly mapping the combined and contrasted signals from specific subsets of cones onto specific behavioural programmes. At this stage of retinal processing, the incoming ‘spectral layering’ is translated into a ‘behavioural layering’^9^, in the sense that the dendrites of ganglion cells associated with specific behaviours systematically take their inputs from specific subsets of inner retinal layers. For example, in zebrafish, ganglion cells associated with the processing of widefield motion vision systematically stratify in the outermost layers of the inner retina, while ganglion cells associated with prey capture stratify in the centre^9,28,31,63^. Interestingly, similar behavioural layering is conserved in mammals^51^, despite their greatly reduced capacity for spectral layering^49,64^.

Together, in zebrafish the systematic inner retinal mapping of different cone mixtures provides a simple and flexible bridge from cones to behaviour, while the simultaneous presence of On and Off signals within individual inner retinal layers allows for local balancing of driving versus regulatory inputs. In support, the retinal output to the brain of fish^46^, birds^65^, reptiles^61,66^ and amphibians^56^, but not mammals^51^, is dominated by On-Off circuits^22^. Any ‘missing’ cone mixtures, including presumably important balancing contributions, can then be provided by amacrine cells^48,67^.

## METHODS

### Animals

All procedures were performed in accordance with the U.K. Animals (Scientific Procedures) Act 1986 and approved by the animal welfare committee of the University of Sussex. Adults and larval zebrafish were maintained at 28°C on a 14:10 hour light:dark cycle. Embryos and larvae were raised in fish water. For all experiments, we used 6-8 days post fertilization (dpf) zebrafish (*Danio rerio*) larvae. For behavioural assays we used wild-type (AB) or *nacre -/-* zebrafish. The latter are mutants without pigments in the skin, but they retain wild-type eye pigmentation. For imaging recordings, 0.1 mM 1-phenyl-2-thiourea (PTU; Sigma-Aldrich, P7629) has been added to fish water from 1 dpf to prevent melanogenesis. The mounting procedure for *in vivo* two-photon imaging was described previously^71^. Briefly, zebrafish larvae were embedded in 1.5% w/v low gelling temperature agarose (Sigma-Aldrich, A9414), placed on a microscope slide oriented dorsal side up and submerged in fish water. To prevent movement, α-bungarotoxin (1 nL of 2 mg/mL; Tocris, catalog no. 2133) was injected into the ocular muscles behind the eye. For all experiments in this study the following previously published transgenic lines were used: *Tg(elavl3:H2B-GCamP6s)*^72^, *Tg(opn1sw1:nfsBmCherry)*^73^*, Tg(opn1sw2:nfsBmCherry)*^74^, *Tg(trβ2:Gal4; UAS:nfsBmCherry)*^75^. The previously unpublished *Tg(LCR:nfsBmCherry)* line was provided by Takeshi Yoshimatsu. Outcrosses of the above transgenic lines were performed to co-express the nuclear-localized calcium indicator GCaMP6s at a pan-neuronal level and the nfsB gene in each cone type. Embryos positive for the transgenes obtained from these outcrosses were isolated and raised to adulthood following standard procedures.

### Two-photon calcium imaging and visual stimulation

Imaging during visual stimulation was performed with a Movable Objective Microscope (MOM)-type 2P microscope [designed by W. Denk, Max Planck Institute (MPI), Martinsried; purchased through Sutter Instruments/Science Products] equipped with a mode-locked Ti:Sa laser (Chameleon Vision-S, Coherent) tuned to 920 nm for GCaMP6s excitation and a water immersion objective (W Plan-Apochromat 20x/1.0 DIC M27, Zeiss). For image acquisition, we used custom-written software [ScanM, by M. Mueller (MPI, Martinsried) and T. Euler] running under IGOR Pro 6.3 for Windows (WaveMetrics). Zebrafish brains were imaged at 1.95 Hz (256x256 pixels, 2 ms per line, 0.87 µm/pixel). For visual stimulation, we custom-built a four-colour hyperspectral spatial stimulator based on a previous design^23^ with new custom software (see below). The four LEDs used were as follows (from ‘red’ to UV): B5B-434-TY, SMB1N-D470-02, SMB1N-420H-02 (all from Roithner), and LZ1-00UV0R (LuxiGen). The LEDs were equipped with the following band- pass filters: FF01-586/20 (Semrock), ET480/40x (Chroma), ET420/40m (Chroma), and FF01-370/36 (Semrock), respectively, to restrict the LED emission spectrum to a narrow band for selective cone excitation. Effective LED spectral peaks as measured at the sample were 587, 470, 422, and 373 nm, respectively. To time-separate scanning and stimulating epochs, LEDs were synchronized with the scan retrace at a line-rate of 500 Hz. Each LED’s intensity was measured and adjusted to follow the relative distribution of the four wavelength peaks of daytime light in the zebrafish natural habitat (from ‘red’ to UV: 1200, 600, 300, and 150 µW/cm^2^). The stimulus sequence described below was projected onto a custom screen based on 63gsm tracing paper (3 cm wide and 1.9 cm high, 1.6 cm from the fish) and presented monocularly. Visual stimuli were executed using the Python software QDSpy (RRID:SCR_01698, see below) as follows: static gratings (9.5° wide, 5 s), gratings moving backwards (9.5° wide moving at 46 °/s, 5 s), static gratings (5 s), dark screen (‘Off’, 10 s), light dots (three bright dots, 4.6° each, bouncing on top of the dark screen in random trajectories at 46 °/s, 5 s), Off (5 s), bright screen (‘On’, 10 s), dark dots (three dark dots bouncing on top of the bright screen, 5 s), On (10 s). Total stimulus duration was 55 s, and this sequence was presented 6 times. To limit the influence of time-dependent effects, the first repeat was excluded from analysis.

### QDSpy

QDSpy is an open-source software package for visual stimulation^23^ that runs under Python3 (https://github.com/eulerlab/QDSpy). QDSpy stimuli are written as normal Python scripts that utilize functions from the ‘QDS’ package to define stimulus objects, set colours, start movies, send trigger signals, and control presentation and display settings (for details, see Ref^23^). For the current study, the software was adapted to present spatial visual stimuli with up to 6 independent ‘colours’ channels simultaneously. To this end, it controls two DLP (digital-light processing) projectors (‘LightCrafter’, DLPLCR4500EVM, Texas Instruments) equipped with custom LEDs via a light guide (see above). The projectors’ optical paths were combined with a dichroic mirror (T400LP, Chroma, F79-100) to co-project their images onto the screen. One projector generated the RGB, the other the UV stimulus components. For stimulus presentation in dual-display configuration (‘screen overlay mode’), the software opens a window that spans both LightCrafter displays and draws the stimuli separately with the appropriate colours onto the two window halves. To ensure that the projector’s built-in video circuits do not change image colours or produce scaling artifacts, the projectors were run in ‘pattern mode’ (which allows exact control over image generation and bit depth) and at the DMD’s native spatial resolution (912 x 1,140 pixels).

### Two-photon data analysis

#### Preprocessing, ROIs, and quality filtering

Recordings were detrended and regions of interest (ROIs) were placed as described previously^46,52^. In brief, we used cell-lab^52^ to segment all somata within a recording plane, and in parallel computed a response-correlation projection of the entire stack, where each pixel is correlated with each of its immediate neighbours to estimate their response coherence over time^46^. Only ROIs that fell within a single soma-sized segment and that passed a minimum response coherence threshold of 0.1 were considered for further processing. We then extracted each ROI’s brightness over time and z-normalised each based on their baseline activity 5 s before stimulus presentation. To limit the effect of time-dependent effects, the first (of 6) response loop was discarded. From here, we optionally applied a combination of response and quality criteria for different analysis, as detailed individually below. First, we computed a response quality index (QI) following Ref^52^, which for each ROI and all of its repeats quantifies the mean of the response variance divided by the variance of the response mean. The resulting index varies between 0 (perfectly random) and 1 (all responses identical), and we used QI>0.5 for inclusion. Second, we computed relative response amplitudes to individual stimulus segments relative to a ROI’s peak response to any stimulus aspect (=1). Only responses with response amplitudes (RA) >0.5 were included. Where relevant, response amplitudes were additionally related across different ‘colour’ stimuli in the same way (i.e. RA = 1 denotes the peak response to any stimulus aspect in any colour). Third, we used the spatial location of ROIs within the brain. For this, we defined the centre of the contralateral or ipsilateral tectum (position 0,0), and only included ROIs within a radius of 80 pixels (69.6 μm).

*‘Population averages’* (e.g. Fig. 2e) were computed based on QI>0.5 without further amplitude filtering, but only for ROIs within the spatial definition of the contralateral or ipsilateral tectum (see above), as indicated. Traces shown simply reflect the sums of all ROIs that pass the above criteria.

*‘Brain maps’* (e.g. Fig. 2f) were computed based on QI>0.5 and RA>0.5 without spatial filtering. Maps were computed based on the original pixel grid of the recordings (i.e. 256x256). For each included ROI across all fish, we added its RA to its spatial location. For better visualisation, we applied x3 spatial binning followed by a 3-pixel SD box smooth to each map.

*‘Trace heatmaps’* (e.g. Fig. 2g) depict all extracted ROI’s z-normalised average response traces within the specified spatial regions of the brain. No QI or RA filtering was applied. ROIs were sorted first by their responses to gratings and bright dots and their relative amplitudes (top), followed by dark-dot dominant responses (middle) and then Off and On-dominant responses, as indicated (bottom).

‘*Response Boxplots across colours*’ (e.g. Fig. 2h) are based on QI>0.5 and RA>0.5 and shown for contralateral ROIs only. In each case, plots show the number of ROIs responding to a given stimulus aspect as indicated (e.g. gratings) relative to the total number of ROIs included.

‘*Responses per fish*’ (e.g. Fig. 3a) denote the total number of contralateral ROIs with QI>0.5 during white stimulation, minus the corresponding number of ipsilateral responses.

*‘Response percentages’* (e.g. Fig. 3e) shows the number of contralateral minus ipsilateral ROIs with QI>0.5 and RA>0.5 per fish, normalised across all five stimuli (=100%). Note that this representation accentuates small response numbers (e.g. see red ablation). For non-normalised brain wide number of responses see each corresponding supplemental panel.

‘*ROI-wise response amplitudes across colours*’ (e.g. Fig. 5a) are based on contralateral ROIs across all fish, with QI>0.5 with no amplitude filtering. Each ROI’s responses were amplitude-normalised to 1 across all colours.

‘*Across-fish correlation’* (Fig. 6a) was computed based the distribution of ‘response percentages’ (see above) across the five stimulus aspects (i.e. gratings, bright dots etc.). For controls and each ablation condition, we computed the correlation coefficient between the distribution of its five response aspects of each fish against the mean of all other fish. Accordingly, a perfectly homogeneous response distribution across the population of fish would yield a correlation of 1, while a perfectly random population would yield 0.

### Cone ablations

To selectively ablate *nfsB*-expressing cones, zebrafish larvae were screened for the expression of the co-expressed fluorescent protein mCherry in the eye and treated with Metronidazole (Met; Sigma, M3761) as described previously^73^. In brief, cone ablation was induced at 5 *dpf* by immersing larvae in fish water containing 10 mM Met for 2 hours for acute UV cones ablation; 5 mM Met for 1 hour for acute ablation of red and blue cones; 10 mM Met for 1h for acute green cones ablation. Following Met treatment, zebrafish were transferred into fish water without Met and fed regularly until used for 2P imaging and behavioural experiments at 6-8 *dpf*.

### DNA extraction and genotyping

Pairs of primers were designed based on the plasmids used to generate the nfsB-expressing transgenic lines, to selectively amplify different DNA sections simultaneously. These custom primers were used in the polymerase chain reaction (PCR) to isolate larvae expressing the nfsB gene in multiple cone types, subsequently employed in double and triple cones ablation experiments. The sequences of each pair of primers were as follows: sw1:nfsBmCherry (forward 5’-TCAAGAAACTCGTGAGGGGT-3’ and reverse 5’-TCAACAACCAGCTTCAGCCA-3’, product length 773 bp); sws2:nfsBmCherry (forward 5’-GCTGGTGACAACAAACCTCA and reverse 5’-GTGCGAGGCATCAAGCATTT-3’, product length 552 bp); LCR:nfsBmCherry (forward 5’-GCAAATGTCCTAAATGAATTTGTGT-3’ and reverse 5’-ATAAAATGCCACGGCTGGGA-3’, product length 435 bp); trβ2:Gal4 UAS:nfsBmCherry (forward 5’- TGCGCCAAGTGTCTGAAGAA-3’ and reverse 5’-TCCGATGATGATGTCGCACT-3’, product length 320 bp). Outcrosses between adult zebrafish expressing Tg(elavl3:H2B-GCaMP6s; opn1sw2:nfsBmCherry) and Tg(elavl3:H2B-GCaMP6s; LCR:nfsBmCherry) were performed. The embryos obtained by this outcross positive for the transgenes (judged by the co-expression of GCaMP in the brain and mCherry in the eye) were isolated and raised to adulthood. Once adult, a genomic DNA extraction was performed to isolate fish expressing the nfsB gene in both green and blue cones. Genomic DNA extraction from adult zebrafish was performed by caudal fin clipping^76^. Animals were anesthetized in Tricaine 25X (4 g/L; Sigma, A5040) and the caudal fin was removed using a scalpel blade. The tissue was then transferred into a reaction tube (Eppendorf) containing 100 µL of 50 mM NaOH (Sigma, 221465), heated to 95°C for 20 min and then cooled to 4°C. To neutralize the basic solution 10 µl of 1 M Tris-HCl, pH 8.0 was added (1/10th of initial volume) followed by a centrifugation at 12000 rcf, 25°C for 5 min, so that the supernatant was ready for use in PCR. To select fish for double cone ablation we used multiplexed PCR, thus amplifying different DNA sequences in one reaction using multiple pairs of primers. For 25 µL reaction with two primers pairs the following reagents were used: 0.5 µL of 10 µM forward primer_1, 0.5 µL of 10 µM reverse primer_1, 0.5 µL of 10 µM forward primer_2, 0.5 µL of 10 µM reverse primer_2, 4 µL DNA sample, 12.5 µL OneTaq Quick-Load 2X Master Mix with Standard Buffer (Biolabs, M0486S), 6.5 µL Nuclease-free water. Reactions were amplified using the following conditions: 94°C for 30 s; 30 cycles of 94° for 30 s, 58°C for 1 min, 68°C for 1 min; followed by 58°C for 5 min. 7 µL of each reaction were loaded onto a 2% agarose gel prepared mixing 2 g agarose (Meridian bioscience, BIO-41025),100 mL TAE 1X (Tris-Acetate-EDTA, pH 8), 2.5 µL SYBR Safe DNA Gel Stain (S33102, Invitrogen) and electrophoresed at 90V for 40 min in TAE 1X. We also loaded 2 µL low range DNA ladder (Meridian bioscience, BIO-33056). The gel was imaged using a ultraviolet (UV) transilluminator apparatus (Odyssey XF, Licor). The bands matching the target size of 435 bp and 552 bp indicated the presence of nfsB-mCherry construct in green and blue cones, respectively. These fish were then incrossed, screened and raised to adulthood and subsequently outcrossed with wild-type fish and genotyped to isolate the homozygous for *nfsB*-mCherry in green and/or blue cones.

For the other cone ablation combinations (e.g. red/UV double ablation, red/green double ablation, green/UV double ablation, red/green/blue triple ablation, green/blue/UV triple ablation) we followed Wilkinson and colleagues’ method for genotyping of live larval zebrafish^77^. Positive larvae obtained by the outcrosses of the transgenic lines for the above-mentioned cone ablation combinations were screened (judged by the co-expression of GCaMP in the whole brain and mCherry in the eye) and isolated for genotyping at 2 *dpf*. Larvae were anesthetized in Tricaine 25X, placed under a stereomicroscope, and the tip of the caudal fin was removed using a 20g needle. The larvae were then transferred in Petri dishes with fish water and kept at 28°C to allow caudal fin regeneration and their normal development. The fin tissue was transferred into a reaction tube (Eppendorf) containing 50 µL of 50 mM NaOH (Sigma, 221465) using a glass pipette, heated to 95°C for 10 min and then cooled to 4°C. To neutralize the basic solution 5 µl of 1 M Tris-HCl, pH 8.0 was added followed by a centrifugation at 12000 rcf, 25°C for 5 min. A multiplexed PCR was performed as described above, but in this case 6 µl of DNA sample were added to the reactions (instead of 4 µl) as well as two or three pairs of primers based on the cone ablation combinations needed. Reactions were amplified and electrophoresed as described above. The bands matching the target size of 320 bp, 435 bp, 552 bp and 773 bp indicated the presence of nfsB-mCherry in red, green, blue and UV cones, respectively. The samples expressing nfsB-mCherry in two and three cone photoreceptors were used for 2P imaging and behavioural experiments at 6-8 *dpf* after Met treatment at 5 *dpf*. Larvae not used for experiments were raised to adulthood.

### Indoor Optomotor assay and spontaneous swimming

Behavioural assays were performed on individual larvae (6-8 *dpf*) using a closed loop optomotor setup as described previously^35^. Freely-swimming larvae were placed in a 6 cm diameter watch glass in filtered fish water. Swimming behaviour was recorded at 200 Hz using a high-speed camera (Omron Sentech STC-CMB200PCL-NIR, Alrad Instruments Ltd, UK) connected to a digital frame grabber (Euresys Grablink Full XR, Stemmer Imaging, Germany) equipped with a zoom lens (Thorlabs, MVL8M23) and infrared (IR) filter (Thorlabs, FELH0750). The arena was illuminated by an IR LED (Thorlabs, M850L3) from below. Stimuli were projected onto a diffuser screen from beneath after reflection by a 5 cm diameter cold mirror (Thorlabs, FM203) using a commercial DLP projector (VAMVO Ultra Mini Portable DLP Projector, 1920x1080 pixels, Shenzhen, China). All experiments were performed in the dark at 28°C. Animals were allowed to adapt to the arena and light conditions for at least 15 minutes prior to each behavioural test. One larva at a time was tested and the same fish were used both for spontaneous and OMR behaviour. For spontaneous swimming we tracked fish both in the dark and in white (RGB) light, 3 minutes each. For optomotor behaviour, image processing and stimulus generation were performed in Bonsai using the BonZeb package^78,79^. Widefield sinusoidal gratings of 1 cm spatial frequency moving at 1 cm/s were presented for 30 s. A closed-loop assay was performed where the orientation of the drifting gratings (calculated using the heading angle of the fish) updated in real time running parallel relative to the fish body axis. Each trial started with a 30 s grey screen, followed by 60 s of gratings drifting rightwards and leftwards (30 s per direction). This protocol was repeated three times, and each session lasted about 5 min. Each recording was then down sampled to 50 Hz, and all subsequent analysis was performed using custom-written Igor Pro 9 scripts (WaveMetrics). Bouts and turns were detected by thresholding each fish’s movement and turn traces over time, with thresholds of 3 pixels/s (∼180 μm / s) and 0.2 radians /s respectively. The exact choice of these thresholds did not qualitatively affect the results. Shown are either the total number of bouts or turns across the full recording (i.e. 3 times 30 s = 90 s), or the corresponding numbers normalised by those of control animals, as indicated. The optomotor index was computed from across both motion directions (see above), as (C-I)/(C+I), where C and I denote the number of correct (towards stimulus) and incorrect turns, respectively.

### Outdoor optomotor assay

Zebrafish larvae (*6-8 dpf)* were placed in a custom-built behavioural rig placed outside in the sun (UK weather permitting, cf. Fig. 8g and Fig. S7a). The was based around an open-top 45 L glass aquarium (50x30x30 cm). To limit reflections and to restrict illumination to overhead only, the four sides of the aquarium were covered with black panels (Thorlabs, TB5). A fifth black panel with a rectangular opening for stimulus display (35x15 cm) was placed beneath. The aquarium was filled with fish water to 25 cm depth, and the temperature was kept at 29°C using a 50W heater. Freely-swimming larvae were placed in a 9 cm Petri dish in fish water. Specifically, the dish was placed in a 3D printed holder in the middle of the aquarium, centered above the rectangular hole (Fig S7b) and half submerged in water, ensuring that the distance between the fish and the stimuli was 25 cm. A 3D printed separator was used to split the Petri dish in half allowing to simultaneously record controls and green/blue double ablated fish (genotyped at 2 dpf as described above), about 25 fish per side. Newly positioned animals were allowed to acclimatise for 15 mins prior to experiments, and 10 mins between trials. Stimuli were presented from below using a custom-built motorized conveyor belt, based on 3D printed parts and 1.5x1.5 cm rails (MakerBeam). For the stimuli, light and dark stripes (10° when viewed from a distance of 25 cm) were produced by gluing LeeFilters (#787) strips onto an A3 white paper. Spectra of both light and dark bars were measured using a spectrometer (Thorlabs, CCS200, Fig. S7d). Stimuli were presented by moving the bars at a speed of 20°/s using the Nema17 stepper motor (driven by l298 H-bridge), connected to a 10 kΩ potentiometer and 9V 1A battery, all controlled by an Arduino Uno. An open-loop assay was performed by moving the bars for 30 s in one direction and 30 s in the opposite direction for 3 times (see also Supplemental Video V3). Optomotor behaviour was recorded in both clear and tinted water. The latter was realized by adding food dyes (red and blue, spectra resultant spectra shown in Fig. S7d). These spectral distortions were introduced to approximate different naturalistic water conditions – note that despite their very different overall shapes, spectra were still wide and without gaps. Behavioural variance attributable to different water conditions was below the natural variance across experimental days and batches of fish (shown separately in Supplemental Figs. 7e,f).

Animals were filmed 10 Hz using a camera (Basler, ACa1440-200um USB 3.0 camera, 1.6 MP) positioned above and equipped with an objective (Fujinon, DV3-4x3-8SA-1 F1.4 f3.8-13mm 1/2") and infrared filter (850 nm, MaD Cameras). Additional infrared illumination (beyond the light provided by the sun) was presented from above using an LED panel to limit shadows provide a homogeneous illumination. Animal tracking was performed in Fiji. Fish positions were used to calculate a running Preference Index (PI) as the difference between the number of fish in the right and left half of the Petri dish divided by the total number of fish (Fig. 8h). From here, we then quantified the ‘population optomotor response’ (population OMR) as the mean position of all fish in the dish over time, corrected for stimulus direction and excluding each first 15 seconds following stimulus reversals. In parallel, we also computed a population heterogeneity index by quantifying the mean squared error between each individual experimental run and the mean of all other runs (such that 0 indicates zero variation across the population).

### Phototaxis

Zebrafish larvae (6-8 *dpf*) were tested for phototaxis in a custom-built behavioural rig, with ten freely-swimming larvae placed in a 5 cm diameter Petri dish in filtered fish water. The Petri dish was placed on a custom 3D printed holder with a separator in the middle to allow the illumination only in one half of the dish. Animals were tracked at 30 Hz using a camera (Basler, ACa1300-200um USB 3.0 camera, 1.3 MP) positioned above and equipped with an objective (Basler, Ricoh Lens FL-CC0614A-2M F1.4 f6mm 2/3") and an IR filter (760 nm, Zomei). Infrared illumination and light of different wavelengths were projected from below onto a diffusing screen. For stimulation we used four LEDs (from ‘red’: Thorlabs, LED591E; Roithner, RLS-5B475-S, VL415-5-15, XSL-365-5E) located both on the left and the right side below the dish so that they were properly collimated and aligned covering the entire half-dish. All the LEDs were controlled using an Arduino Mega with custom scripts. The intensity of each LED was adjusted to 6 µW/cm^2^ on the sample. All experiments were performed in the dark and at a controlled temperature of 28°C. Animals were allowed to adapt to the arena and light conditions for at least 15 minutes before starting the behavioural tests. Visual stimuli were presented alternately on one side of the dish for 30 s, three times per side (each session lasted 3 min, see also Supplemental Video V4). We waited at least 5 minutes prior to the presentation of the next stimulus (i.e. different wavelength light). Each recording was then down sampled to 1 Hz and multi-animal tracking was performed using the Fiji *mosaic* plugin. Fish positions were used to calculate a running Preference Index (PI) as the difference between the number of fish in the lit side and the dark side divided by the total number of fish. From here, we calculated a phototaxis index per dish (of 10 fish) and trial based on the average PI between 10 and 30 seconds into each stimulus switch (i.e. leaving out the first 10 seconds, while fish were mostly still moving). The exact choice of this time window did not qualitatively affect the results. Analyses were performed with custom-written MATLAB and Igor Pro 9 scripts.

### Natural scene processing

(Fig. 1) A luminance-calibrated RGB underwater video of the zebrafish natural habitat based on Ref^70^ where a robotic gantry system was used to gradually advance the camera through vegetation over a distance of ∼ 30 cm was used. Water depth was ∼50 cm. For processing, brightness was calculated as the sum of the three spectral channels, while ‘spectral width’ (‘whiteness’) was computed per frame and pixel, as 1 minus the standard deviation divided by the mean across the three spectral channels. In this way, a pixel with equal red, green and blue values yield a ‘whiteness’ of 1 (independent of brightness), while unequal values across the three channels yield correspondingly lower values. Notably, there are many ways that could be used to estimate spectral width from RGB video data, but because these all yield qualitatively similar results, we opted to use this very simple metric above.

Activation of opponent and non-opponent channels as a function of decreasing whiteness (Fig. 1b-d) was computed based on the simplifying assumption that full-spectrum illumination is spectrally sinusoidal from 0-π (cf. Fig 1d), gradually narrowing along a triangle formed from 0 and π towards a monochromatic point at 0.75 * π. ‘Whiteness’ (x-axis in Fig. 1c). is then computed as the full width half maximum of this model spectrum. Spectra were then multiplied with a same frequency and phase-aligned sin^2^ function to mimic non-opponent drive (Fig. 1d, bottom), and a correspondingly frequency-doubled sin^2^ function for opponent drive (Fig. 1d, middle). Normalised activation was then computed as the area under the curve relative to zero in each case, divided by the largest entry across whiteness conditions in each case.

### Modelling of cone drive

(Fig. 5). To estimate the underlying cone contributions that led to the spectral tuning functions of grating responses and bright dot responses following green/blue double ablation and in controls, we set up a cone-combinatorial model. Red and UV cone contributions were fitted to the tuning functions from green/blue ablated animals, while independently the contributions from green and blue cones were estimated based on the differences in spectral tunings functions between controls and green-blue double ablated animals. Except for these spectral target functions, and the cone pairs used to fit them, the remainder of the modelling was identical, and will therefore only be described once, using the example of red/UV fitting for gratings (Fig. 5c) as an example.

We used the previously determined^13^ spectral tuning functions of the cones as the input (shown in Fig. S1b, and converted to expected activations for the stimulus combinations in this present work in Fig. S1d,e). First, we computed the sum of both cones’ tunings for all possible cone ratios (i.e. r*red + u*UV, where r and u each varied between −1 and 1.), and mapped them onto a circle indicated as ‘linear’ around the origin of the heatmaps shown, such that r;u = 1;0 and r;u = 0;1 were mapped to the right and top of the circle, respectively. The four quadrants therefore represent all possible positive sums (top right), negative sums (bottom left) or opponent contributions (top left, bottom right), while the axes represent pure cone contributions (e.g. right is +red only, top is +UV only, and so on). We then computed corresponding nonlinear versions of the same basis functions by raising them to the power of *nl*, which ranged from 0 to 5. Accordingly, *nl*<1 gave sublinear scaling, while *ln*>1 gave in supralinear scaling. This nonlinearity *nl* was mapped along the radial dimension as indicated. Based on this spatial mapping, we then shaded each pixel by the mean squared error (MSE) between each resulting template function and the target tuning function as defined by the corresponding brain responses (cf. Fig. S4e-h). MSEs were normalised to the error score resulting from a fit with all entries at zero, such that MSE = 0 indicates a perfect fit, while MSE =>1 indicates that the fit was equal than or worse than no fit at all. In each case, the fit with the lowest MSE overall (filled symbol) and the lowest MSE of a linear model (open symbol) are indicated.

### Quantification and statistical analysis

No statistical methods were used to predetermine sample size. Owing to the exploratory nature of our study, we did not use randomization or blinding. We used 1 and 2-tailed paired and non-paired Wilcoxon Rank Sum or T-Tests with Bonferroni correction for multiple comparisons for all statistics, as appropriate and individually indicated.

## ACKNOWLEDGEMENTS

We thank Takeshi Yoshimatsu for providing the original cone-ablation lines, including the previously unpublished green-cone ablation line, Patricio Simoes and Leon Lagnado for access to their OMR setup, and Emma Alexander for providing the underwater video used in Figure 1. We also thank Michel Laurin for key discussions around of vertebrates’ aquatic histories, and Leon Lagnado for critical feedback on a draft of the manuscript. Funding was provided by the Wellcome Trust (Investigator Award in Science 220277/Z20/Z), the European Research Council (ERC-StG “NeuroVisEco” 677687 and ERC-AdG “Cones4Action” covered under the UK’s EPSRC guarantee scheme EP/Z533981/1), UKRI (BBSRC, BB/R014817/1 and BB/W013509/1), the Leverhulme Trust (PLP-2017-005, RPG-2021-026 and RPG-2-23-042) and the Lister Institute for Preventive Medicine. This research was funded in part by the Wellcome Trust (220277/Z20/Z). To promote Open Access, the authors have applied a CC BY public copyright licence to any Author Accepted Manuscript version arising from this submission.

## AUTHOR CONTRIBUTIONS

see table

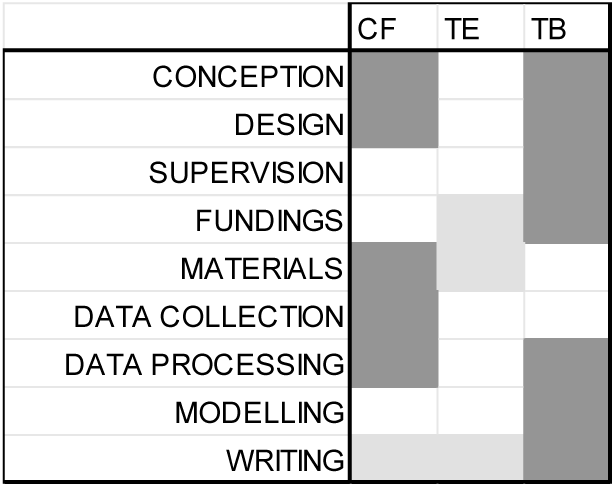

## SUPPLEMENTAL TABLES AND VIDEOS

**Supplemental Tables T1-7.** Explanatory Legends are included in the tables themselves.

**Supplemental Video V1.** Full movie from example underwater scene from the zebrafish natural habitat as shown in Fig. 1a (see also Methods). Note how ‘whiteness’ (right) but not ‘brightness’ systematically demarcates the visual foreground.

**Supplemental Video V2.** Example recording as Fig. 2b from the same optic tectum plane of a 7 dpf control larva during the presentation of the visual stimuli indicated in the upper left corner. Spectral composition of the stimulus (RGBU, RGB, U) is indicated by the squares in each panel. Movie speed x2.

**Supplemental Video V3.** Example recording of outdoor optomotor behaviour (cf. Fig. 8g) in control (top half) and green/blue double ablated (bottom half) larvae. Movie speed x5.

**Supplemental Video V4.** Example recording of phototaxis with red light in wild type larvae (cf. Fig. 8k). The illuminated side is indicated by the red square. Movie speed x4.

## SUPPLEMENTAL FIGURES

**Figure S1 Related to Figure 2.**
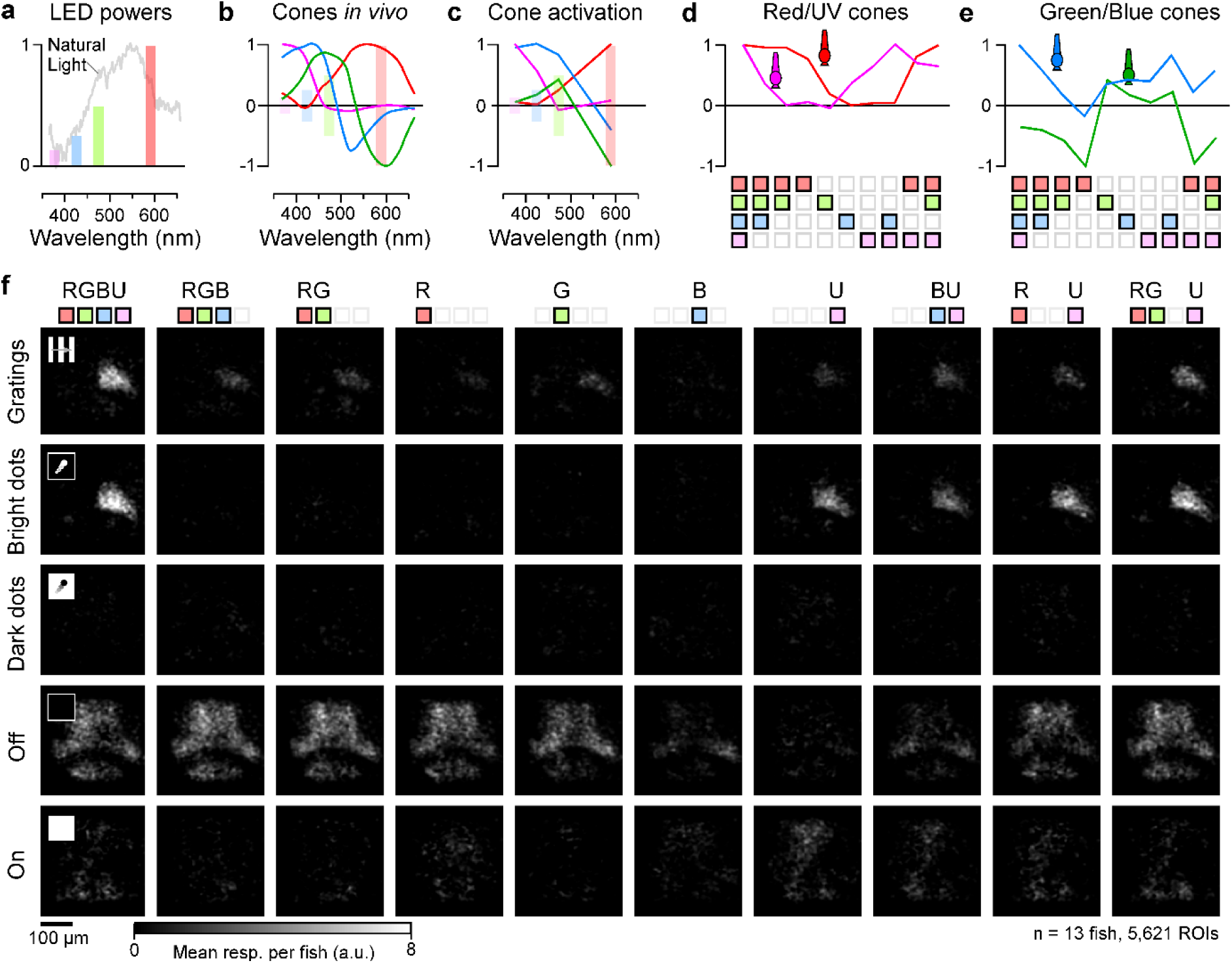
**a**, Spectral centres of light emitting diodes (LEDs) used for visual stimulation including their relative powers, superimposed the mean spectrum of natural daylight in the zebrafish natural habitat (based on Ref^24^). **b-e**, Full spectral tuning functions of larval zebrafish cones in vivo (b, based on Ref^13^) superimposed on LED positions as shown in (a), the same cone’s spectral sensitivities specifically at the LED positions multiplied by the respective LED powers (c), and the corresponding individual cone activation patterns for all presented LED combinations (d, e, cf. Fig. 2). **f**, as Fig. 2f, here shown for all presented LED combinations.

**Figure S2 Related to Figure 3.**
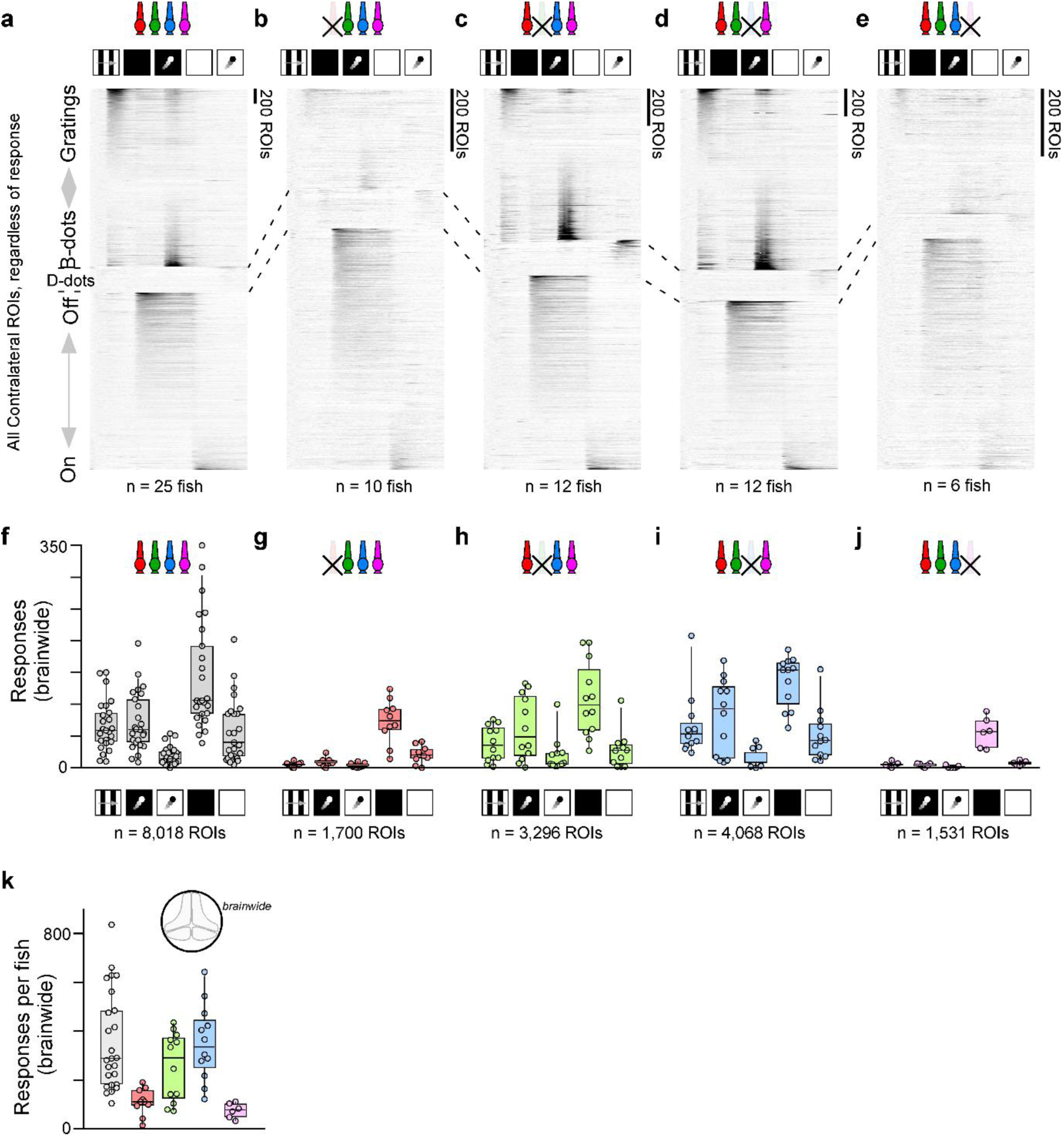
**a-e**, as Fig. 2g, but here shown for white-only stimulation of controls (a) and following ablation of different cone types: red (b), green (c), blue (d) and UV (e). **f-j and k**, as Fig. 3e-I and a, respectively, but here shown for uncorrected numbers of responses brain wide.

**Figure S3 Related to Figure 4.**
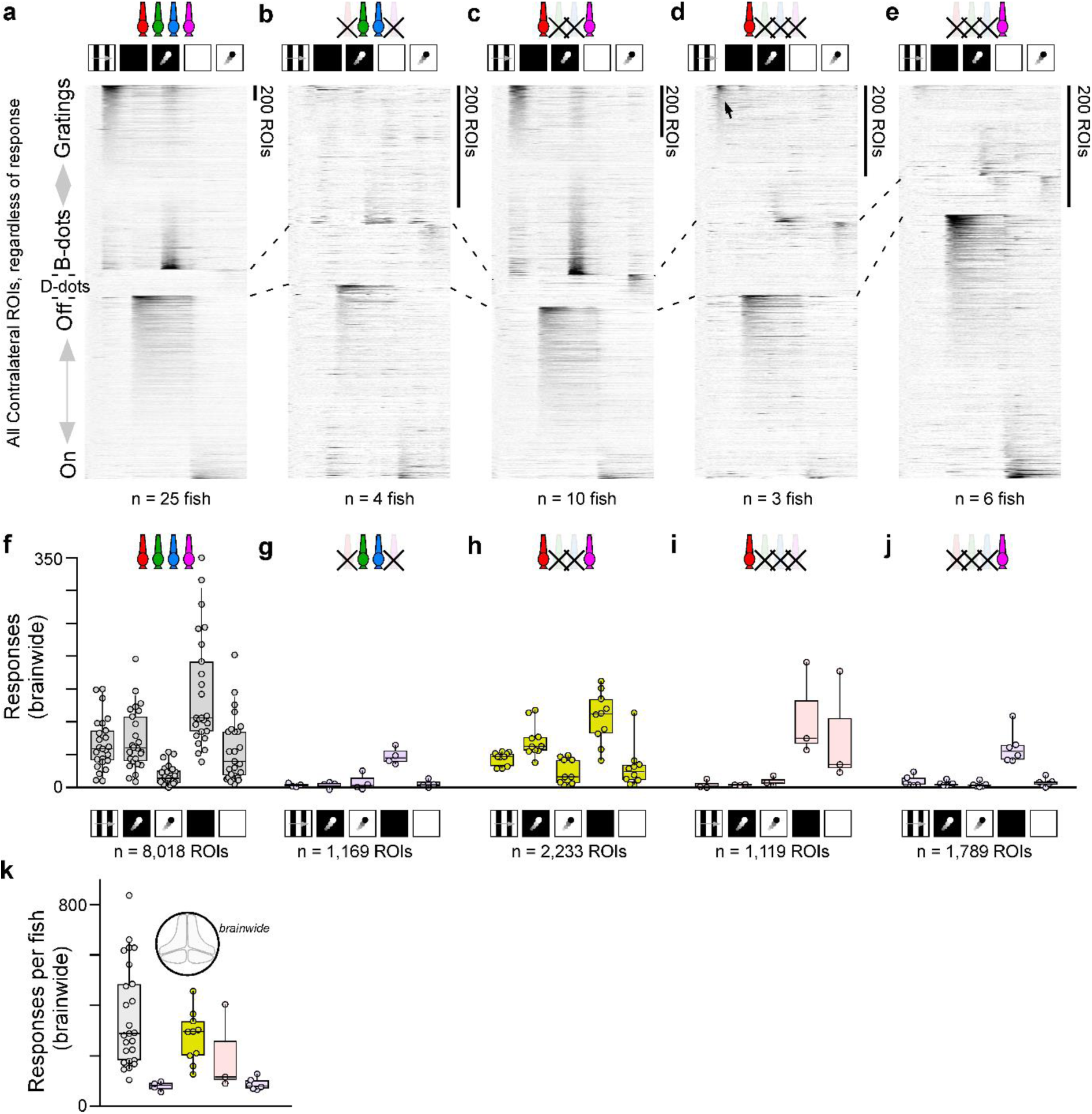
**a-k,** as Supplemental Fig. S2 a-k, but here shown for the different set of cone ablations as shown in Figure 4.

**Figure S4 Related to Figure 5.**
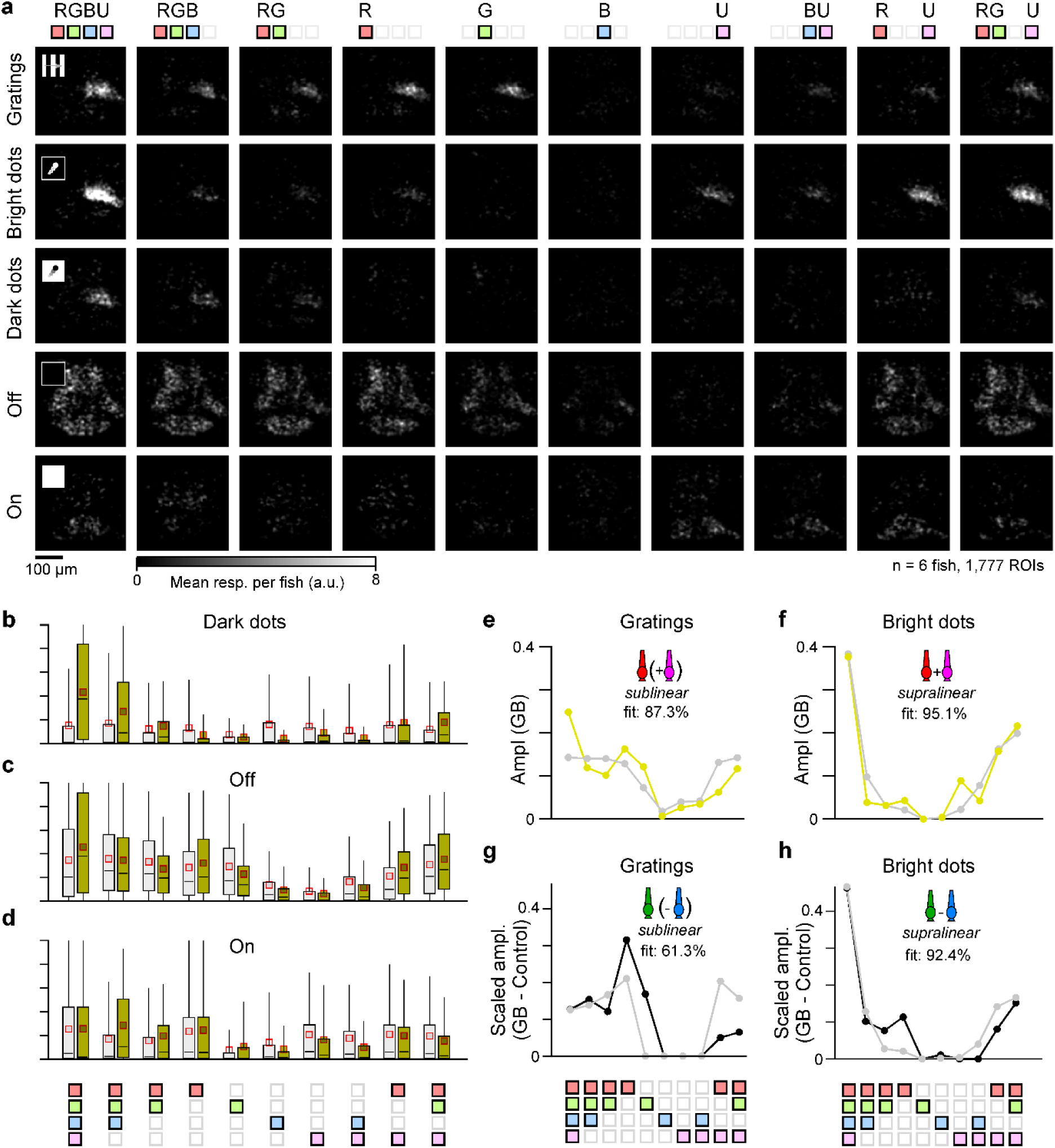
**a**, as Fig. S1f, but here shown for green/blue double ablated animals. **b-d**, as Fig. 5a,b, but here shown for the remaining three stimuli as indicated. **e-h**, related to Fig. 5c-f, corresponding best model fits (grey) superimposed on respective target tuning functions (yellow/black). Indicated fit quality based on the residual mean squared error (MSE) of the best fit as shown, compared to the fit with all entries = 0.

**Figure S5 Related to Figure 6.**
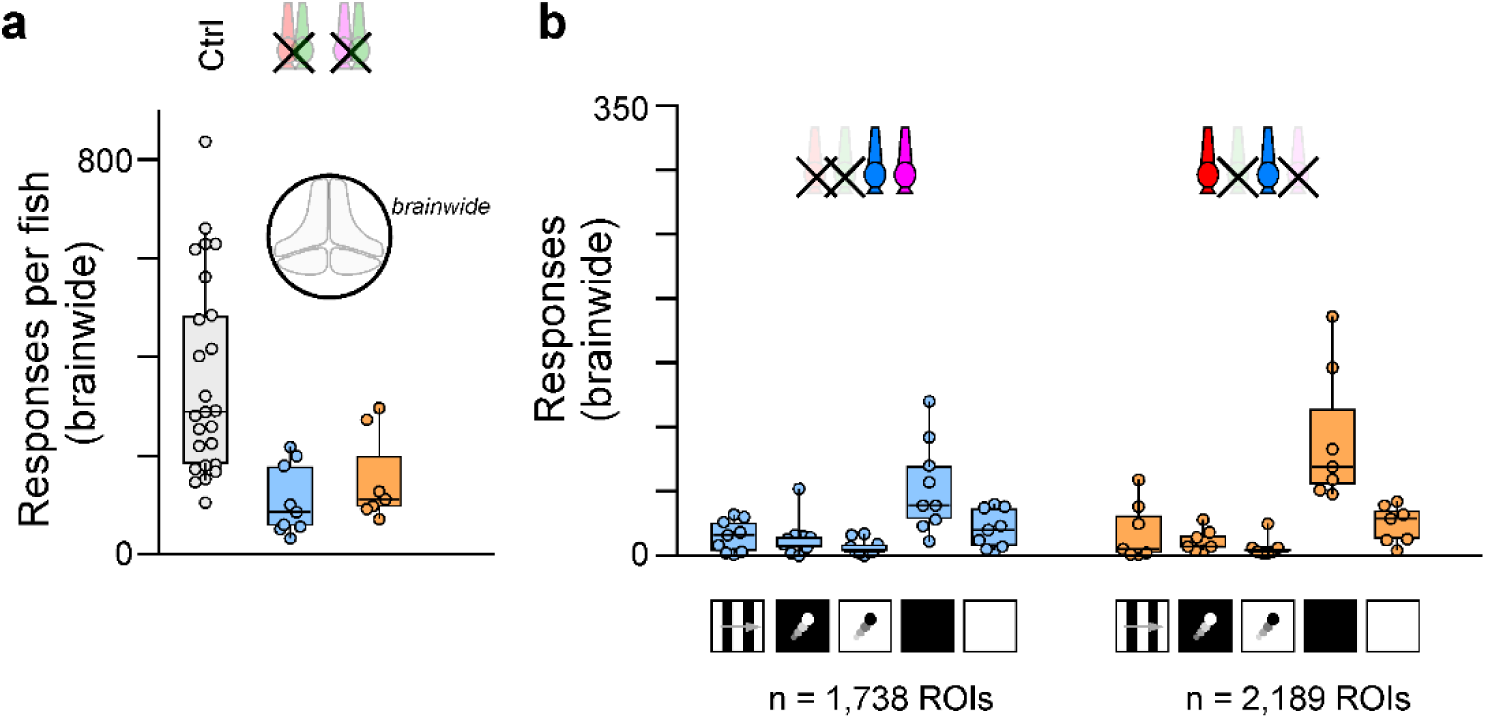
**a,b,** as Fig. S2f,k, but here shown for the new ablation combinations as introduced in Fig. 6.

**Figure S6 Related to Figure 7.**
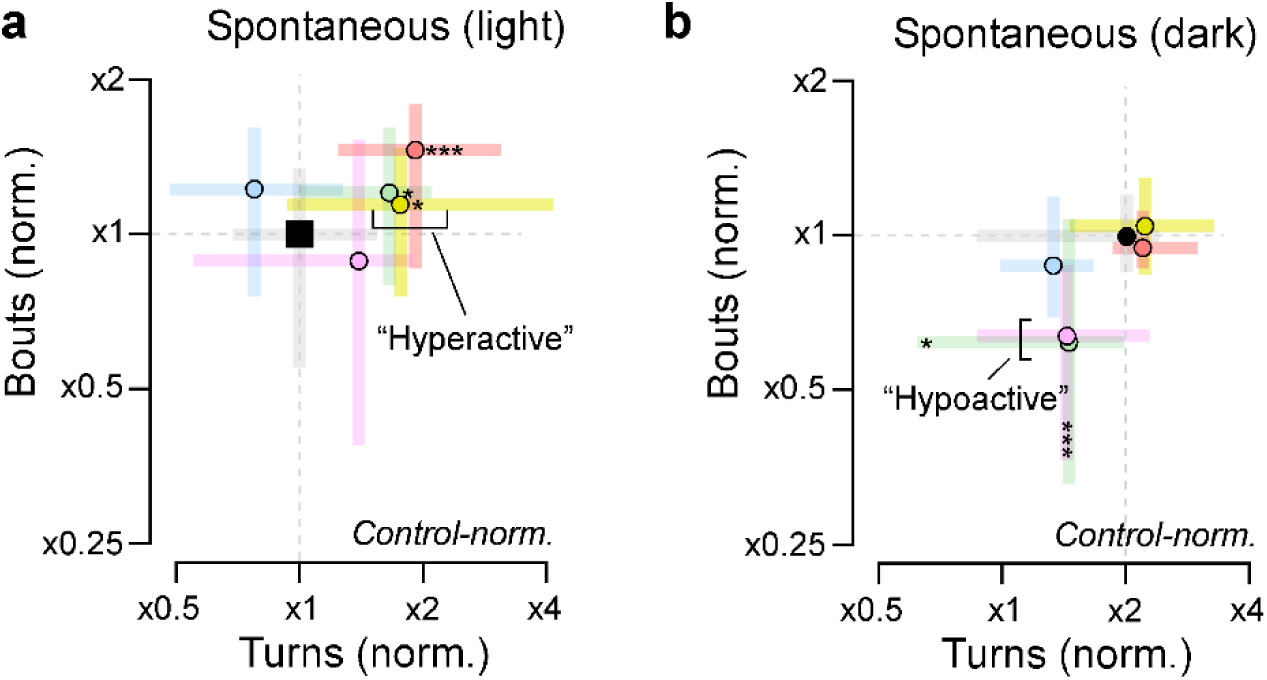
**a,b,** As Fig. 7e,f, but for normalised bout and turn rates in the light (a) and dark (b). For full statistics, see Supplemental Table T5.

**Figure S7 Related to Figure 8.**
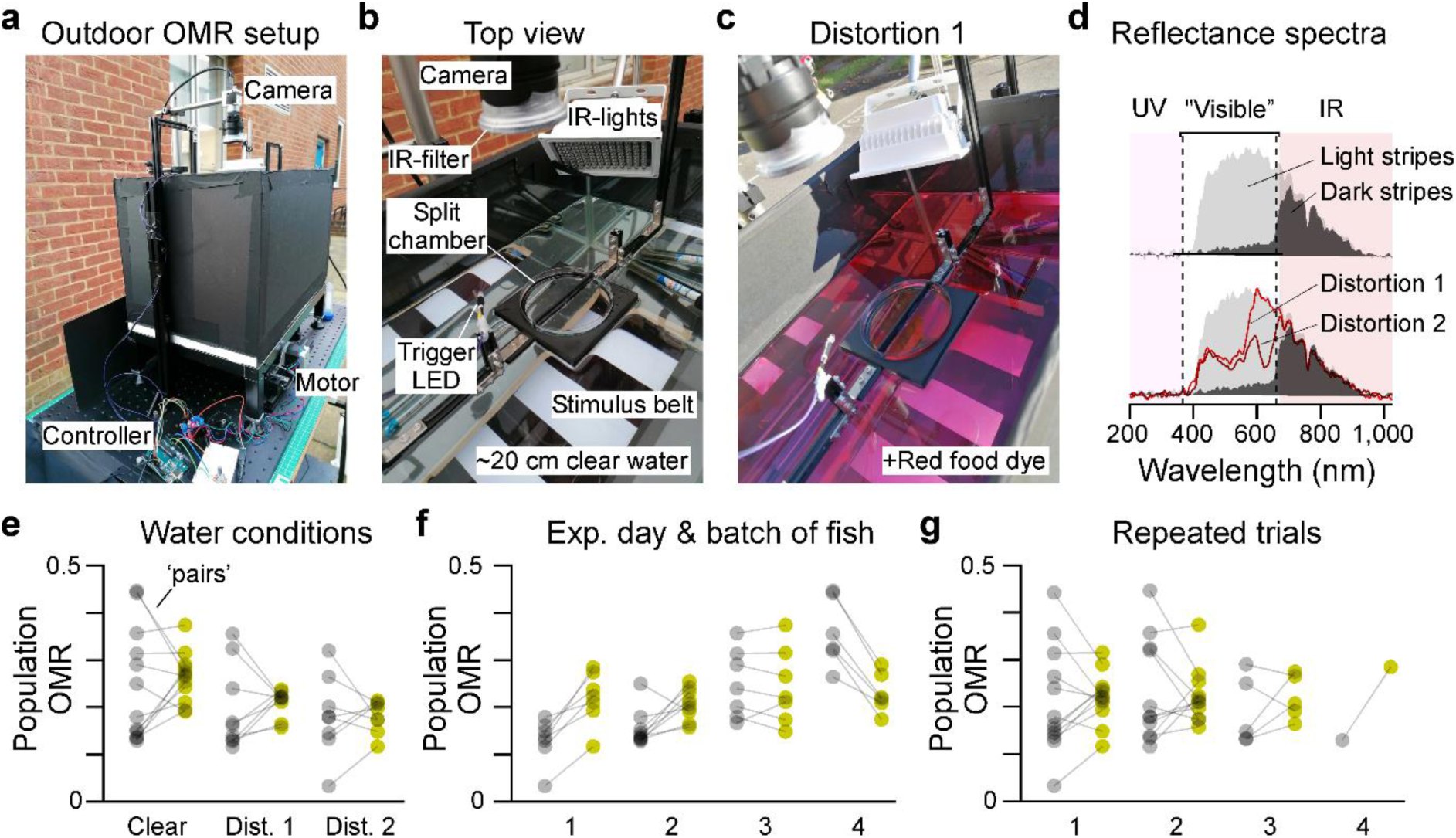
**a-c,** Photographs of the outdoor optomotor setup, showing the full system (a), a top-view close-up (b) and a corresponding close-up after adding ‘red’ food dye to the water (distortion 1, Methods), used to mimic potentially varying turbidities encountered in nature. **d**, Natural sunlight reflectance spectra measured as seen by the fish, for the dark and light stimulus bars as indicated (top) and for the light bars following two tiers of spectral distortions (cf. c and Methods). Note that despite the distortions, all spectra remained broad from UV to red, and without notable spectral gaps. **e-f**, Optomotor performance sorted by water condition (e, clear water and two tiers of distortions), experimental day / batch of fish (f, different batches of fish were used on different days), and repeated trials for individual batches and conditions (g).

## SUPPLEMENTAL DISCUSSION

### Ancestral blue cones in platypus?

Platypus represent the only possible exception to the complete absence of ancestral blue cones in mammals, because their genome uniquely retains the ancestral blue opsin gene (SWS2^41^). However, opsin identity is an incomplete predictor of cone identity, because cones routinely alter opsin expression (discussed in Ref^9^). In fact, platypus lack the ancestral UV opsin (SWS1), potentially hinting that they, like all other mammals, exclusively retain ancestral red and UV cones, with UV cones expressing SWS2 opsin.

### Revisiting Walls’ nocturnal bottleneck theory

In mammals, the loss of ancestral green and blue cones has traditionally been linked to Walls’ ‘nocturnal bottleneck theory’. This long-standing interpretation start with the assumption that ancestral blue and green cones mainly or exclusively serve colour vision and posits that therefore their retention in the mammalian eye provided little competitive advantage during the age of the dinosaurs, when mammals occupied nocturnal niches for some 180 million years. However, it may be time to revisit this long-standing idea.

First, an assumed lack of exploitable spectral information at night as a leading driver for cone loss is questionable. ‘Colour information’ (better: ‘spectral diversity’) does not disappear at night: To a first approximation, nocturnal skylight is spectrally identical to daylight, because it is dominated by the sun’s reflection in the moon. Correspondingly, ‘nocturnal colour vision’ is readily achieved by boosting the sensitivity of ‘daylight photoreceptors’, and this has independently evolved in multiple lineages of both vertebrates (e.g. amphibians, geckos, snakes) and invertebrates (e.g. some moths or beetles).

Second, as argued in the main text, factors other than colour vision might have driven the mammalian loss of ancestral green and blue cones. In zebrafish, one of these functions relates to underwater foreground enhancement, but it seems unlikely that this is their only ‘non-colour’ function. Here, better understanding what other factors might have contributed to their loss will require continued exploration of their functions in the eyes of diverse vertebrates that retain them.

Third, and perhaps most importantly, the loss of mammals’ ancestral blue/green system does not plausibly coincide with the start of the nocturnal bottleneck around 243 million years ago. First, ancestral green cones could have been lost as early as 330 million years ago, preceding the first dinosaurs by 87 million years. Second, if as argued (see above), platypus indeed lack ancestral blue cones, then the same is also true for this second auxiliary cone type. However, even if platypus does indeed retain ancestral blue cones, the timelines do not work: In this case, blue cone loss probably occurred after the divergence of monotremes (platypus) and marsupials (187 mya), but before the divergence of marsupials and eutherian mammals (147 mya). This is because no marsupial or eutherian mammal retains ancestral blue cones, so the most likely timepoint for blue cone loss is before their divergence. Accordingly, in this scenario, ancestral blue cones would have had to persist a minimum of 44 million years of continuous nocturnalisation in the lineage that led to all extant mammals, and indeed for the full 180 million years of nocturnal bottleneck in the lineage that led to modern day platypus.

### Why two ‘regulatory’ cones?

The use of two rather than one type of colour-opponent cone to regulate red/UV circuits brings about at least two possible benefits for underwater vision.

First, it adds flexibility. Beyond ‘whiteness’, behaviourally relevant stimuli differ in their spectral properties. For example, larval zebrafish preferentially use the long-wavelength biased^24^ riverbed as a visual anchor for optomotor behaviours^69,70^, while their zooplanktonic food when illuminated by the sun is short-wavelength biased^29^. Here, the use of two rather than one regulatory cone system may add important flexibility to usefully segment both types of signals.

Second, it adds robustness: Variations in illumination, water depth and/or dissolved solutes can all shift in the average spectral content of light in aquatic habitats^10^. Accordingly, the use of two opponent circuits with distinct spectral zero crossings^13^ increases the probability that at least one of them robustly tracks visual distance. Moreover, wiring the two systems in mutual opposition presumably allows detecting deviations from ‘white’ in two spectral directions, and/or for both negative and positive contrasts.

